# Profiling the bloodstream form and procyclic form *Trypanosoma brucei* cell cycle using single cell transcriptomics

**DOI:** 10.1101/2023.01.09.523263

**Authors:** Emma M. Briggs, Catarina A. Marques, Guy R. Oldrieve, Jihua Hu, Thomas D. Otto, Keith R. Matthews

## Abstract

African trypanosomes proliferate as bloodstream forms and procyclic forms in the mammal and tsetse fly midgut, respectively. This allows them to colonise the host environment upon infection and ensure life cycle progression. Yet, understanding of the mechanisms that regulate and drive the cell replication cycle of these forms is limited. Using single cell transcriptomics on unsynchronised cell populations, we have obtained high resolution cell cycle regulated transcriptomes of both procyclic and slender bloodstream form *Trypanosoma brucei* without prior cell sorting or synchronisation. Additionally, we describe an efficient freeze-thawing protocol that allows single cell transcriptomic analysis of cryopreserved *T. brucei*. Computational reconstruction of the cell cycle using periodic pseudotime inference allowed the dynamic expression patterns of cycling genes to be profiled for both life cycle forms. Comparative analyses identify a core cycling transcriptome highly conserved between forms, as well as several genes where transcript levels dynamics are form-specific. Comparing transcript expression patterns with protein abundance revealed that the majority of genes with periodic cycling transcript and protein levels exhibit a relative delay between peak transcript and protein expression. This work reveals novel detail of the cell cycle regulated transcriptomes of both forms, which are available for further interrogation via an interactive webtool.

## Introduction

The *Trypanosoma brucei* life cycle involves developmental transitions between replicative and cell cycle arrested forms, the latter of which are primed for transmission between the mammal and tsetse fly, or vice versa^1^. Metacyclic trypomastigotes emerge from the replicating epimastigote population as arrested G0 forms in the tsetse fly salivary gland and express genes required for infection of the mammal, which occurs during their transfer during a bloodmeal^2^. Metacyclics subsequently re-enter the cell cycle and differentiate into replicative slender bloodstream forms (BSFs). Slender BSFs proliferate and increase in parasitaemia before exiting the cell cycle via a quorum sensing mechanism^3,4^ and differentiating into G0 arrested stumpy BSFs, which express genes required for differentiation into replicating procyclic forms (PCFs) once in the tsetse fly midgut^5^.

The cell cycle of *T. brucei* broadly follows the typical eukaryotic progression through G1, S, G2 and M phases followed by cytokinesis, although trypanosomes are unusual in that both nuclear and mitochondrial genome replication and segregation are precisely orchestrated during cell division. while many canonical regulators remain unidentified, are absent, or have been replaced by trypanosomatid-specific factors. Several regulators have been identified^6–9^ including cdc2-related kinases^10^ (CRKs) and 13 cyclins^6–9,11^, several of which have been linked to regulation of the *T. brucei* cell cycle phase transitions. Additionally, transcriptomic, proteomic and phosphoproteomic analysis of semi-synchronized PCF populations have uncovered numerous cell cycle regulated (CCR) genes for further investigation^12–14^. However, little overlap has been observed between these studies^14^, reflecting both the variation between experimental design and differences between transcript and protein regulation.

Single cell transcriptomics (scRNA-seq) allows the transcriptomes of individual cells in a heterogenous, asynchronous population to be captured without the need to first isolate the target cell types by methods such as physical or chemical synchronisation and cell sorting. Continuous biological processes, such as cell cycle progression and cellular differentiation, can then be reconstructed *in silico* using trajectory inference and pseudotime approaches where cells are ordered by their progressive transcriptomic changes^15,16^. Differential expression analysis across these ordered cells identifies genes with altered transcript levels during the process, and the dynamic change in transcript levels can be modelled.

scRNA-seq has been used effectively to compare transcriptomes of various *T. brucei* life cycle stage forms, including those extracted from tsetse flies to analyse the development of metacyclics in the salivary gland^17–19^ and to investigate slender to stumpy differentiation of BSFs *in vitro*^20^. These studies mainly employed droplet based methods (Drop-seq^17^ and Chromium 10x Genomics^18,20^) to recover higher cell numbers and relied on live, freshly derived parasites to ensure sufficient transcript recovery per cell^21^. However, the need to use live parasites for droplet-based methods restricts usage of these approaches in experiments where high cell numbers or multiple time points are required, for example when modelling a developmental processes with trajectory inference methods^15^. A previous attempt to use methanol fixed BSFs with Chromium technology yielded low transcript recovery per cell^21^.

In this study, we profile the dynamic transcript changes during the cell cycle of laboratory cultured ‘monomorphic’ slender BSFs (refractory to stumpy differentiation and so quorum sensing dependent cell cycle exit) and PCFs. For each form, we also compare 10X Chromium generated transcriptomes from live parasites and parasites cryopreserved with glycerol in liquid nitrogen (LN_2_). We find cryopreservation causes limited changes to the transcriptome of BSF and PCF *T. brucei*, highlighting cryopreservation as a valuable method of sample preservation for scRNA-seq analysis of trypanosomes. This will allow for future studies involving multiple conditions, or sampling over a time course using trypanosomes and, likely, other kinetoplastida and apicomplexan parasites. Periodic pseudotime inference was applied to the resulting data to model the cell cycle progression of both BSF and PCF *T. brucei*, allowing the genes with CCR transcripts to be identified in each form. Comparison with existing high quality PCF proteomic datasets^13,14^ further revealed a relative offset in peak transcript and protein levels for at least 50% of genes exhibiting CCR with respect to transcripts and proteins. Comparison between BSFs and PCFs identified genes with shared or life cycle stage-specific CCR transcripts, revealing both common and developmentally-specific CCR factors, as well as apparent differences in the S-G2 transition between forms.

## Results

### Cryopreservation of *T. brucei* for Chromium single cell transcriptomics

Generating scRNA-seq data with droplet-based technology Chromium (10x Genomics) currently requires live trypanosome samples in order to recover a high number of transcripts per cell^21^. To test whether *T. brucei* could be stored prior to processing, we compared the impact of cryopreservation using 10% glycerol on live cell recovery when using a slow thawing protocol (Fig. S1, methods). Using motility as a measure of parasite viability indicated both BSF and PCF cells recovered with high viability after freezing with 10% glycerol, with each form maintaining at least 90% cellular motility after 28 days of cryostorage (Fig. S1a). When returned to culture, parasites showed a delayed return to normal growth rates (Fig. S1b, c) indicating samples should be processed for scRNA-seq immediately after thawing to reflect their transcript status when cryopreserved.

Using this approach, replicating BSF and PCF *T. brucei* were processed for Chromium scRNA-seq “fresh” from *in vitro* culture or after 13 days of storage with 10% in LN_2_, hereafter referred to as “frozen” (Fig. S2). Frozen samples were thawed on day 13 and processed alongside the fresh samples taken directly from culture, thus fresh and frozen samples contain biological replicates and were subjected to scRNA-seq in the same batch.

Cryopreservation had little effect on the raw data quality (Fig. 1a, b, c; supplementary data 1) with the total numbers of unique transcript counts (unique molecular identifiers; UMIs) and features (encoding genes) detected per cell unaffected for either BSF or PCFs (Fig. 1a, b).

**Figure 1.**
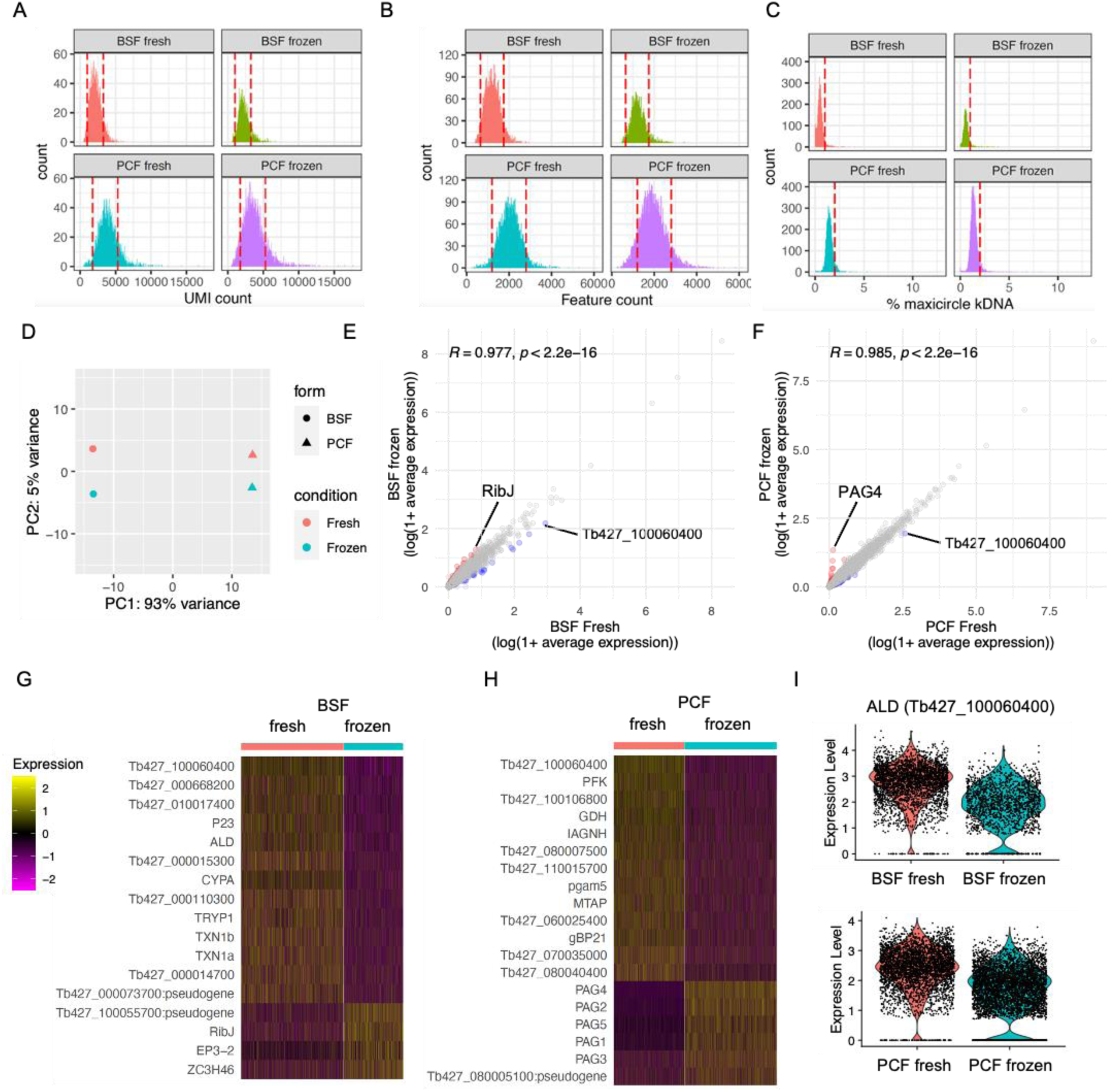
scRNA-seq of cryopreserved and fresh *T. brucei* BSF and PCF. **a**) The unique molecular identifiers (UMI, x-axis) captured per cell (count, y-axis) captured by Chromium scRNA-seq with BSF and PCF taken fresh from *in vitro* culture (fresh) or after cryopreservation in LN_2_ (frozen). Red dashed lines indicate threshold used for QC filtering of each sample. **b**) Number of genes (features, x-axis) for which transcripts were capture per cell. **c**) Percentage of transcripts captured per cell that are encoded by genes on the mitochondrial maxicircle kDNA genome (% maxicircle kDNA, y-axis). **d**) Top two components (PC1 and PC2) identified with PC analysis after pseudobulking all counts for each sample. Fresh (red) and frozen (blue) samples are shown for BSFs (circle) and PCFs (triangles). **e**) Average expression of each gene across all cells for BSF fresh (x-axis) and BSF frozen (y-axis) plotted as log(1+mean average count). Correlation coefficient and p-value of one-tailed wilcox test is indicated above. Gene with increased FC > 2 in frozen sample are coloured red and those decreased in blue. **f**) Average gene expression of PCF samples, as in e. **g**) Scaled expression of genes DE between fresh and frozen BSF scRNA-seq (adjusted p-value < 0.05, FC > 1.5). Gene names are given when available, otherwise gene IDs are shown. **h**) as in g for PCF samples. **i**) Raw transcript counts (expression level) for fructose-bisphosphate aldolase (ALD; Tb427_100060400) in BSF (upper) and PCF (lower).

Additionally, the percentage of transcripts derived from the mitochondrial kDNA maxicircle genome was unchanged by the freezing and recovery procedures (Fig. 1c). The percentage of kDNA-derived transcripts was higher in PCF compared to BSF, as expected: only PCFs require complexes III and IV for oxidative phosphorylation^22^, components for which are encoded on the kDNA maxicircle^23^. Higher average UMIs and features per cell were also recovered in PCF compared to BSF, in both fresh and frozen samples, although it is unclear if this is a biological phenomenon or if RNA extraction and capture is more efficient from PCFs. After filtering the transcriptomes based on these parameters to remove those of low quality or likely multiplets (Fig. 1a-c), 81.7% and 81.04% of fresh and frozen BSFs cells were retained in the data leaving 2,767 and 1,599 total cells, respectively. For PCFs, 76.82% and 72.60% of fresh and frozen cells were retained, leaving 3,305 and 4,335 cells respectively.The differences in total number of cells are likely due to variation in loading and cell capture between samples.

Principle component analysis (PCA) highlighted far great variability between samples of different life cycle forms (93% of variance), compared to the preparation method (i.e. fresh or frozen, 5% variance) (Fig. 1d). Average transcript counts across cells for each gene were significantly correlated between fresh and frozen samples in both BSF and PCF forms (Pearson’s *R* = 0.977 and 0.985, respectively) (Fig. 1e, f), with few genes (BSF: 0.80% of genes captured, PCF: 0.55%) showing greater than 2-fold difference (supplementary data 2). DE analysis comparing single cell transcriptomes of fresh and frozen samples using MAST^24^ revealed 17 genes altered in BSF (14 upregulated in fresh, 3 in frozen) and 19 genes (13 in fresh and 6 in frozen) between PCFs (adjusted p-value < 0.05, FC > 1.5) (Fig. 1g, h; supplementary data 2). Only one gene, which putatively encodes fructose-bisphosphate aldolase class-I, was differentially expressed in both forms, with higher expression in fresh samples (Fig. 1i). Notably, procyclin associated genes (PAGs) 1-5 were all upregulated in frozen PCFs (Fig. 1h).

As no large-scale transcriptomic changes in response to cryopreservation were observed, and DE genes did not include those linked to cell cycle regulation, fresh and frozen samples were integrated as replicate samples to analyse the cell cycle of PCF and BSF *T. brucei*.

### The cell cycle regulated transcriptome of PCF *T. brucei*

PCF scRNA-seq data from fresh and frozen samples were integrated and dimensional reduction was performed. Transcriptomes were then plotted in low dimensional space as unifold manifold approximation and projection (UMAP^25^) plots, where cells are arranged by transcriptional similarities and differences (Fig. 2a). Using cell cycle phase markers identified previously using bulk-RNA-seq^12^, each cell was labelled by phase (Fig. 2b). Grouping by phase was evident in both samples, with each population arranging in a logical order according to cell cycle progression. The proportion of cells in each phase was similar between samples (Fig. 2c) and corresponded with the proportion of cells G1 (1N), S (>1N <2N) and G2/M (2N) phases as assessed by flow cytometry analysis of DNA content (Fig 2d). A proportion of cells (fresh 6.23%, frozen 12.02%) did not elevate transcript levels of markers for any phase and so were named “unlabelled” (grey; Fig 2b, c). The majority of unlabelled cells cluster with early G1 cells (Fig. 2b). DE analysis between early G1 and unlabelled cells found 14 genes with adjusted p-value <0.05, yet none showed fold-change >1.5 (supplementary data 3). These include three ribosomal proteins, a DEAD box helicase and a putative subunit of replicative protein A (RPA). It is possible that these cells are yet to re-enter the cell cycle and so do not over express any early G1 markers, or that the early G1 markers used here are insufficient to label all cells in this phase.

**Figure 2.**
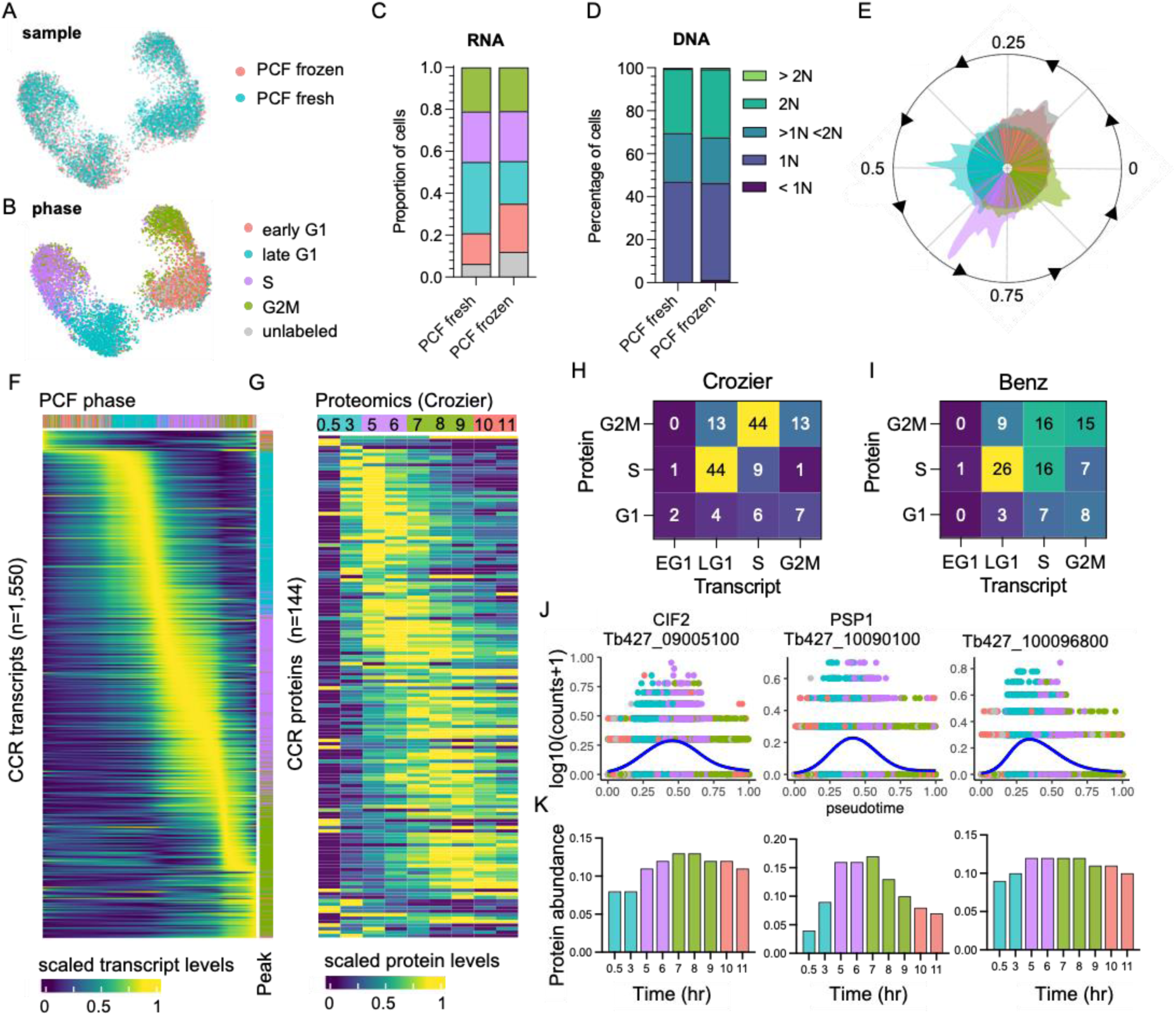
The cell cycle transcriptome of PCF *T. brucei*. (**a**) UMAP plot of integrated PCF transcriptomes from fresh (blue) and frozen (red) samples. (**b**) UMAP of PCF transcriptomes coloured by inferred cell cycle phase. (**c**) Proportion of cells assigned to each phase by transcriptomics as in b. Legend as in b. (**d**) Proportion of cells with DNA content assessed by flow cytometry. (**e**) Histogram of transcriptomes arranged in pseudotime (anti-clockwise) representing cell cycle progression. Each line in inner circle indicates one transcriptome coloured by phase as in **b**. Outer circle histogram of showing number of cells at each point in pseudotime (0-1). (**f**) Scaled transcript levels of CCR genes (rows), ordered by peak time, plotted across transcriptomes (columns) ordered in pseudotime. Top annotation indicated cell phase, right annotation indicated phase with highest expression of each gene. (**g**) Scaled protein abundance for 129 genes identified as CCR by Crozier *et a*l, plotted in the same order as f. Time points are indicated in top annotation, coloured by the most enriched cell cycle phase for each sample. Numbers of genes with highest transcript expression in each phase analysed by scRNA-seq (x-axis) and highest protein level identified by Crozer *et al* (**h**) and Benz *et al* (**i**) proteomics studies. (**j**) Transcript counts of three genes (y-axis) plotted across pseudotime (x-axis). Each dots shows one transcriptome coloured by phase as in b. Blue line shows smoothed expression across pseudotime. (**k**) Protein abundance for the same genes as in j, previously identify as CCR by Crozier *et al*. Time point and colour of most enriched phase for each sample (x-axis) as in g.

Pseudotime values were assigned using Cyclum, an autoencoder technique which projects cells on a nonlinear periodic trajectory^26^. This is performed independent of the UMAP plotting and phase assignment described above. Cells ordered according to cell cycle progression and phases clearly separated in pseudotime, with the exception of early G1 and “unlabelled” cells (Fig. 2e). As expected, total RNA increased over pseudotime from early G1 (3,221 median UMI per cell) to G2M (4,077 median UMI) (Fig. S3a). Hence, DE analysis across pseudotime was performed using normalised counts to find CCR transcripts independent of total RNA increase. PseudotimeDE^27^ was used to identify DE genes and thresholds for selecting significant CCR genes were selected based on the detection of the previously identified CCR genes in PCF transcriptomic analysis^12^ (Fig. S4). Using these cut-offs (FDR adjusted p-value < 0.01, mean fold-change > 1.5) 1,550 significant CCR genes were identified (Supplementary data 4), including 399 of the 530 genes (75.28%) previously detected with the bulk RNA-seq approach^12^ (Fig. S5, supplementary data 4). Dynamic expression patterns were evident across the cell cycle (Fig. 2f). Each gene was classified as peaking in a particular phase by comparing the average expression levels across cells for each phase. This revealed 77 (4.53%) genes with highest expression in early G1, 498 (29.31%) in late G1, 598 (35.20%) in S phase, and 526 (31.00%) in G2/M (supplementary data 4, Fig 2f).

### Relative temporal relationship between RNA and protein levels in PCFs

To investigate the correlation between transcript and protein abundance during the PCF cell cycle, CCR genes defined by scRNA-seq above were compared with CCR proteins identified in two separate studies. Crozier *et al*. (2018) and Benz *et al*. (2019) employed centrifugal elutriation to enrich for smaller G1 phase *T. brucei* cells which were then returned to culture and allowed to progress through the cell cycle in a semi-synchronized manner over time. Mass-spectrometry was then employed to analyse protein samples taken as the cell population progressed through the cell cycle. Comparison of protein abundance in each sample then allowed CCR proteins to be identified. 427 and 370 genes (annotated in the WT427 2018 genome) were classified as encoding CCR proteins by Benz and Crozier, respectively, with 61 classed as CCR in both datasets. Of the 1,550 genes with CCR transcripts in the present scRNA-seq data, 226 were classed as having CCR proteins in both or one of these studies (Fig. S5). Proteomics analysis has lower sensitivity compared to transcriptomics, therefore not all scRNA-seq defined CCR genes are detected as proteins in these studies: 998 (64.39%) and 667 (43.03%) CCR genes were detected by Crozier and Benz, respectively. Of these, just 14.43% and 17.69% had been classified as CCR by Crozier (Fig. 2g) and Benz (Fig. S6a), respectively. Thus, the majority of CCR transcripts do not result in CCR protein levels as defined by current methods. Proteins were not detected for 586 scRNA-seq CCR transcripts in either study and so, could not be compared (Supplementary data 4).

Plotting scaled CCR protein levels in the Crozier data (Fig. 2g) and, to a lesser extent, in the Benz data (Fig. S6a) revealed dynamic abundance patterns across the cell cycle that broadly followed the dynamic transcript patterns identified by scRNA-seq. Comparing the relative timing of peak transcript and peak protein levels showed a common trend where transcript levels often peaked in the phase preceding the protein peak (Fig 2h, i). This broad pattern was observed for 66.67% and 47.22% of CCR genes when comparing transcripts to protein levels from Crozier (Fig. 2h) and Benz studies (Fig. 2i), respectively. Comparison to the Benz study indicated more genes peaking in the same phase for transcripts and proteins (37.78%), compared to the Crozier study (19.44%).

Just 24 genes were classified as CCR in scRNA-seq, bulk-RNA-seq^12^ and both proteomic studies (Fig. S5). These include genes with documented roles in the cell cycle: cyclin-dependent kinase CRK3 (Tb427_100054000), cyclin-dependent kinase regulatory subunit CKS1 (Tb427_110183500), cytokinesis initiation factors CIF1 (Tb427_110176500) and CIF2 (Tb427_090085100), and Cohesin subunit SCC3 (Tb427_100064300). Others include three homologues of *S. cerevisiae* Polymerase Suppressor PSP1 (Tb427_100090100, Tb427_110047000 and Tb427_110165900), and six genes encoding hypothetical proteins with no known function (Tb427_040026500, Tb427_040054600, Tb427_080009800, Tb427_100096800, Tb427_100120400 and Tb427_110082700). Transcript levels for these genes were raised (Fig. 2j) prior to protein levels (Fig 2k, S6c).

### The cell cycle regulated transcriptome of BSF *T. brucei*

The same approach was taken to analyse transcript dynamics during the BSF cell cycle. Transcriptomes from both the fresh and frozen samples (Fig. 3a) were arranged in low dimensional space according to phase assigned using bulk RNA-seq defined markers (Fig 3b). Notably, S and G2M BSF cells display less separation in UMAP plots compared (Fig. 3b) to PCFs (Fig. 2b), indicating less distinction between the transcriptomes of these phases for BSFs. As observed in PCFs, early G1 and unlabelled BSF transcriptomes overlapped significantly (Fig 3b). DE analysis between these two phases identified 16 genes (adjusted p-value <0.05), yet none reaching a FC cut-off of >1.5 (Supplementary data 3). Of these, 3 were also DE between early G1 and unlabelled PCFs: a putative ribosomal protein S9/S16 (Tb427_070014300), a hypothetical protein (Tb427_010013900) and a RPA subunit (Tb427_050022800). The proportion of cells in each phase was similar between fresh and frozen samples (Fig. 3c), as well as phases defined by DNA content (Fig. 3d).

**Figure 3.**
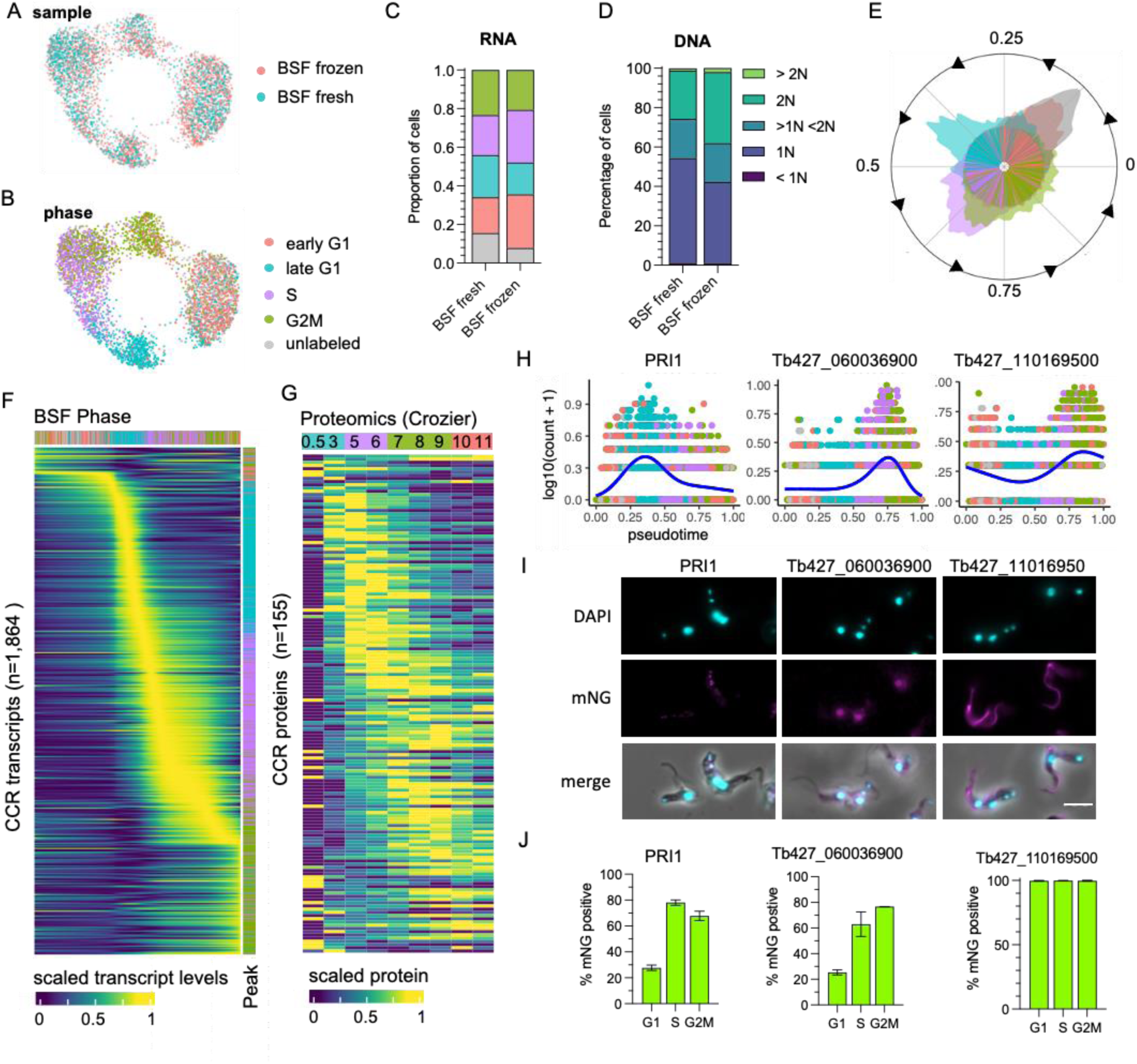
The cell cycle transcriptome of BSF *T. brucei*. (**a**) UMAP plot of integrated BSF transcriptomes from fresh (blue) and frozen (red) samples. (**b**) UMAP of BSF transcriptomes coloured by inferred cell cycle phase. (**c**) Proportion of cells assigned to each phase by transcriptomics as in b. Legend as in b. (**d**) Proportion of cells with DNA content assessed by flow cytometry. (**e**) Histogram of transcriptomes arranged in pseudotime (anti-clockwise) representing cell cycle progression. Each line in inner circle indicated one transcriptome coloured by phase as in **b**. Outer circle histogram of showing number of cells at each point in pseudotime (0-1). (**f**) Scaled transcript levels of CCR genes (rows), ordered by peak time, plotted across transcriptomes (columns) ordered in pseudotime. Top annotation indicated cell phase, right annotation indicated phase with highest expression of each gene. (**g**) Scaled protein abundance for 137 genes identified as CCR by Crozier *et al*, plotted in the same order as f. Time points are indicated in top annotation, coloured by the most enriched cell cycle phase for each sample. (**h**) Transcript counts of three of the top CCR genes (y-axis) plotted across pseudotime (x-axis). Each dots shows one transcriptome coloured by phase as in b. Blue line shows smoothed expression level across pseudotime. (**i**) Fluorescent microscopy imaging of mNeonGreen (mNG) tagged top CCR proteins. DAPI staining of DNA (cyan) and mNG fluorescence (magenta) are shown for the three genes as well as merged with DIC (merge). Scale bar = 10 µm (**j**) The percentage of cells positive for mNG as detected by flow cytometry analysis. For each gene, counts are separated by cell cycle phase, inferred by DNA content detection (G1 = 2C, S = >2C < 4C, G2M = 4C).

Using Cyclum to infer pseudotime during the cell cycle (Fig. 3e), also indicated S and G2M BSF cells were less distinct in their transcriptome than in PCFs at this cell cycle transition. DE analysis over pseudotime identified 1,864 CCR transcripts (FDR adjusted p-value <0.01, FC >1.5) with dynamic expression during the cell cycle (Fig. 3f, supplementary data 5). A remarkably similar proportion of genes peaking in each phase was found for BSF compared to PCF. In BSFs, 76 (4.08%) genes had highest expression in early G1, 588 (31.55%) in late G1, 678 (36.37%) in S phase and 522 (28.00%) in G2M (supplementary data 5, Fig. 3f).

Proteomics data across the cell cycle is not currently available for BSFs; yet, 13.83% (122 of 882 detected) and 12.54% (155 of 1,236 detected) of the BSF CCR transcripts were identified as CCR in PCF proteomics datasets from Benz (Fig S6b) and Crozier (Fig 3g), respectively. The expression pattern of these common CCR genes, largely following the same pattern as PCF, with transcripts peaking prior to protein levels.

To investigate CCR proteins directly in BSFs, the top most significant genes with transcripts peaking in late G1, S and G2M phase were tagged at the N-or C-terminus with the fluorescent epitope tag mNeonGreen (mNG) using CRISPR/Cas9^28^. The top two CCR genes to peak in late G1 were MORN repeat-containing protein 1 (MORN1, Tb427_060051900) and mitochondrial DNA primase, Pri1 (Tb427_080028700) (Fig. 3h). MORN1 has been localised by immunofluorescence previously, revealing the protein is part of the specialised trypanosome bilobe, cytoskeletal structure located close to the flagella pocket^29,30^PRI1 was found previously to locate to the antipodal sites flanking the mitochondrial kDNA in PCFs^31^. Using fluorescence microscopy, we found that tagged mNG::PRI1 (N-terminal tag), akin to the observations in PCFs, also localises to the flanking sides of kDNA in BSFs (Fig. 3i). Flow cytometry analysis was used to compare expression of mNG::PRI1 to cell cycle phase, inferred using DNA content (Fig. 3j and S7c). While 27.71% G1 cells were detected as expressing mNG::PRI1, this increased to 78.06% in S phase cells and 67.73% of G2M phase cells. Additionally, fluorescence intensity peaked in S phase cells (Fig. S7c). Thus, a delay between protein and transcript levels are also notable for PRI1, with transcripts peaking in late G1 (Fig. 3h), but protein in S phase (Fig. 3j and S7c).

The top S phase peaking transcripts were two genes encoding histone H2B (Tb427_100112400 and Tb427_100112200), followed by Tb427_060036900 which encodes a hypothetical protein of no known function (Fig. 3h). N-terminal tagging of this gene was unsuccessful, but C-terminal tagging resulted in viable clones. Fluorescence microscopy revealed nuclear localisation, including in post-mitotic cells where both nuclei contained fluorescent protein (Fig. 3i, S7b). Flow cytometry revealed 25.44% of G1 phase BSFs expressed Tb427_060036900::mNG, increasing to 63.04% and 76.72% of S and G2M phase cells, respectively (Fig. 3j), in keeping with cyclic transcript levels expression increasing in S and G2M phase (Fig. 3h).

The most significant gene peaking in G2M was Tb427_110169500, which also encodes a hypothetical protein (Fig. 3h). N-terminal tagging revealed this protein is expressed in all cycle phases (Fig. 3i, j) and localises to both the old and newly developing flagellum (Fig 3i, S7b). Transcript levels increase as cells progress from into S and peak in G2M, perhaps to meet increased protein requirement as the new flagella develops. Slightly increased fluorescence of mNG was evident for G2M cells compared to G1 (Fig. S7c).

### Common CCR transcripts in BSF and PCF forms

BSF and PCF CCR transcripts were compared to identify 1,013 genes classed as CCR in both forms using a threshold of adjusted p-value <0.01 and FC >1.5 (Fig. S8a-c, Supplementary data 6). Expression patterns appeared to show greater coordination in the early stages of the cell cycle, whereas patterns in the S and G2M phase showed greater variability between forms (Fig. 4a, b). 83.12% (842) of the common CCR genes showed highest transcript levels in the same cell cycle phase, and 16.19% (164) peaked in neighbouring phases (Fig. 4b).

**Figure 4.**
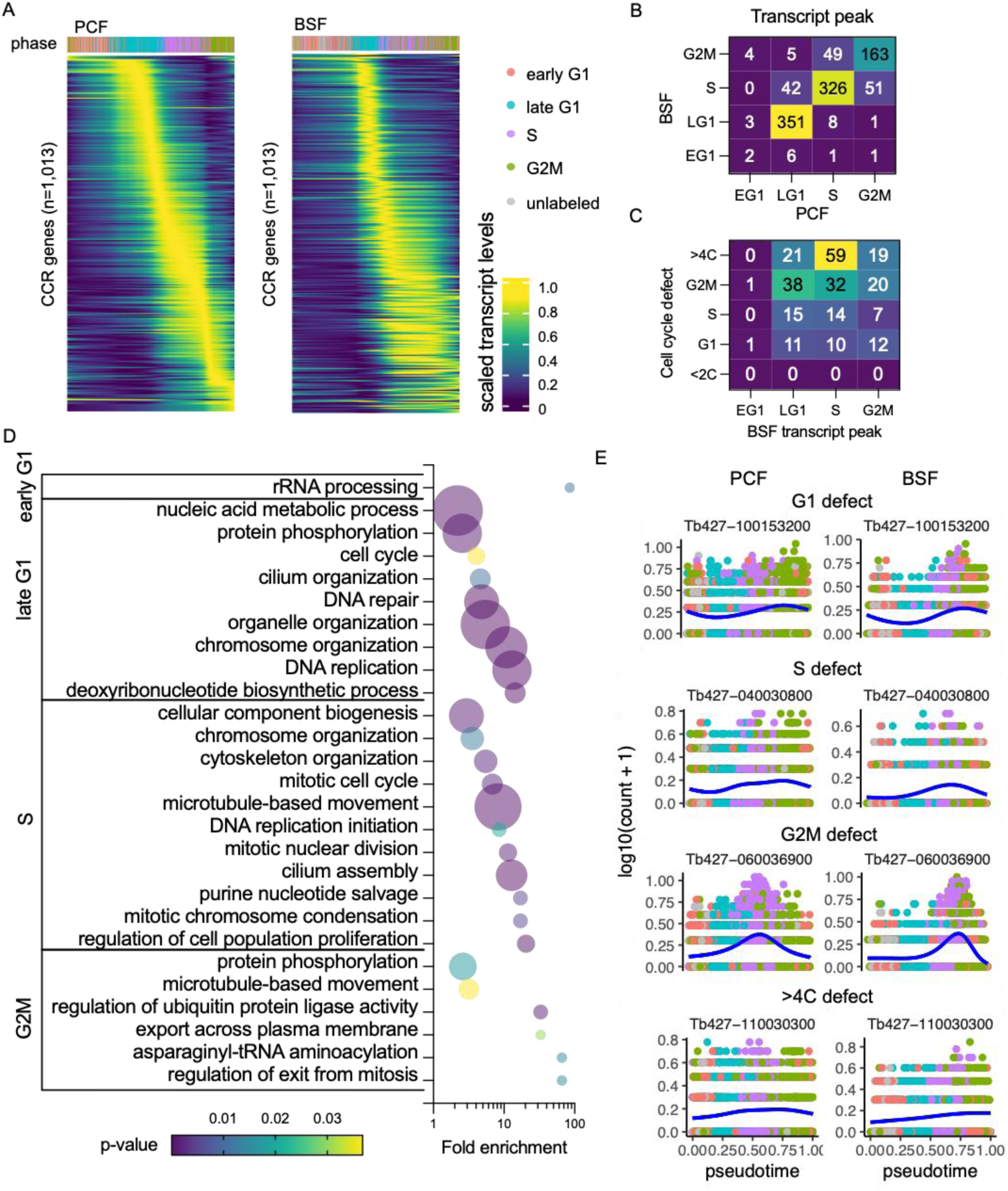
Common CCR transcripts of BSF and PCF *T. brucei*. (**a**) Scaled transcript levels of common CCR genes (rows), ordered by peak time and plotted across transcriptomes (columns) ordered in pseudotime of PCF (left) and BSF (right). Genes are ordered by peak time in PCF in both cases. Top annotation indicates cell cycle phase. (**b**) Number of genes peaking in each cell cycle phase for PCF (x-axis) and BSF (y-axis) transcriptomes. (**c**) Number of genes peaking in each BSF phase (x-axis) linked to a cell cycle defect (y-axis) in RIT-seq screen of BSFs by Marques *et al* 2022. (**d**) GO terms associated with common CCR grouped by peak phase in the BSF cell cycle. Fold change of detected genes is plotted on x-axis, points are sized by the number of genes and coloured by p-value. (**e**) Transcript levels of the most significantly DE gene associated with each cell cycle defect category (G1, S, G2M and >4C). Counts per cell (y-axis) are plotted across PCF (left) and BSF (right) pseudotime (x-axis, coloured by phase as in a. Blue line shows smoothed expression level across pseudotime.

Genes were classified based on the phase with highest expression in the BSF cell cycle (supplementary data 4), and Gene Ontology (GO) analysis was performed to find biological processes associated with each set of genes (Fig. 4d, supplementary data 6). Few GO terms were enriched for early G1 genes, as only 9 genes peak in this phase, and all of these are either labelled as “hypothetical” or have been assigned descriptions based on putative functional domains. All GO terms, including “rRNA processing”, relate to one gene, Tb427_110120100, which shares sequence homology with UTP21, a component of the small-subunit processome^32^.

In late G1, several genes associated with “protein phosphorylation” are evident, including known cell cycle associated genes such as CRK2^10,33,34^, aurora kinase 3 (AUK3^35,36^) and Wee1-like kinase^37^. 14 late G1 genes related to the term “DNA replication”: 5 genes encoding components of the MCM DNA replication licensing complex (MCM2, 4-7), an MCM10 homolog, 5 DNA polymerases, two DNA primase subunits and the DNA synthesis factor RNR1 (ribonucleoside-diphosphate reductase large chain). DNA repair protein RAD9 and recombination helicase proteins PIF1, 2, and 5 show similar regulation, are associated with “DNA repair” and, the in case of the helicase proteins, “telomere maintenance” GO terms.

S phase is associated with GO terms “DNA replication initiation”, due to peak expression of CDC45 (cell division cycle 45) and another MCM component, MCM3. Together with the GINS complex, MCM2-7 and CDC45 form the replicative helicase CMG complex^38^ that is activated only in S phase. Two genes encoding putative components of the condensing complex (CND1 and CND3) are predicted to have roles in “mitotic chromatin condensation” and “mitotic nuclear division”. Mitotic cyclin CYC6 is clearly CCR in both forms and peaks at the S-G2M transition (Fig. S9), while a second mitotic cyclin, CYC8, peaks during late G1 in both life cycle stages (Fig. S9). Kinetoplastid membrane protein 11 (KMP11-2), a known positive regulator of cytokinesis in both BSF and PCF^39^ together with two paralogs, KMP11-1 and KMP11-5, also peaked in S phase.

In G2M phase, AUK1 and AUK2 transcripts are at their highest levels which, along with the kinetochore phosphorylating kinase KKT10^40^ (kinetoplastid kinetochore protein 10) and five other kinases, are enriched for the term “protein phosphorylation”. Six genes associated with “microtubule based process” were upregulated in G2M, including two putative kinesins (one of which, KIN-F, is known to localise to the spindle during mitosis^41^) and KLIF (kinesin localising to the ingress furrow), which is required for cleavage furrow ingression during cytokinesis^42^. CDC20 transcripts are highest in G2M and is associated with the “regulation of ubiquitin protein ligase activity” term; yet, there is no evidence of CDC20 acting on the Anaphase Promoting Complex/Cyclosome (APC/C), at least in PCFs^43^. Other G2M associated genes include a putative homolog of *S. cerevisiae* CDC14, which has several roles in regulating mitosis^44^.

Recently, a genome scale phenotypic genetic screen (RNA Interference Target sequencing, RIT-seq) was performed to identify genes associated with a cell cycle defect when transcripts were depleted by RNAi in BSF *T. brucei*^45^. After induction of RNAi, cells from each cell cycle phase (G1, S and G2M) were isolated based on their DNA content using FACS; sub-diploid (<2C) and over-replication (>4C) populations were also isolated and analysed. Analysis of each pool revealed depletion of which genes had led to an enrichment of parasites in each population, associating a cell cycle defect to 1,198 genes (16.63% of those investigated). Of the 1,013 common CCR genes identified here by scRNA-seq analysis, 260 (25.67%) were shown to have a cell cycle defect using the same threshold as Marques *et al* (Fig 4c, supplementary data 6). The peak transcript expression phase of these genes showed low association with phenotype defect, although 22.69% of genes with highest expression in S phase resulted in a >4C defect, indicating these S phase genes are required for correct genome replication and its control.

As mechanisms of cell cycle regulation and cyclical transcript changes are largely conserved across the eukaryotes, we hypothesised that genes with CCR transcript levels in *T. brucei* are more likely to be conserved. To investigate, we extracted the 819 orthogroups which contained the common CCR regulated genes and compared their conservation across 44 kinetoplastid proteomes, including trypanosome and leishmania species^46^. CCR orthogroups were conserved across significantly (p < 0.0001) more proteomes (mean of 41.66 out of 44 genomes), compared all orthogroups of the *T. brucei* Lister427 reference genome^47^ (mean of 18.44 genomes), and a random subset of 1,000 orthogroups (mean 18.43) (Fig. S8d). More orthologs per orthogroup were also present across the species for CCR genes (mean of 50.79 proteomes) compared to all orthogroup (mean of 23.31 proteomes) and the random subset (mean of 22.45 proteomes) (Fig. S8e). Of the highly conserved common CCR genes, 365 genes are described as encoding “hypothetical” proteins (9.69% of the total hypothetical protein encoding genes located in the core chromosomes), indicating they may have central unknown roles in the kinetoplastida cell replication cycle. Of these, 61 had a cell cycle defect identified by Marques *et al*.; depletion of 9 led to increased BSF in S phase, 23 in G2/M, 14 in G1, and 15 in >4C. The transcript levels for most significant genes for each defect are plotted across the PCF and BSF cell cycle (Fig. 4e).

### Unique CCR transcripts in PCF and BSF

Of the CCR transcripts in PCF, 540 were only significant in this form and showed varied expression across all phases of the cell cycle (Fig. 5a, supplementary data 7). GO term enrichment (Fig 5b, supplementary data 7) of these genes uncovered terms including “lipid metabolic process” attributed to 9 genes encoding putative proteins, including one encoding a putative triacylglycerol lipase which peaks in S phase, and a putative C-14 sterol reductase, for which transcripts are highest in late G1 (Fig. S10a). Genes relating to “ribonucleoprotein complex biogenesis” include ribosome production factor 2 (RPF2), which is part of the 5S ribonucleoprotein (RNP) complex in PCFs^48^, and 20S-pre-rRNA D-site endonuclease, NOB1, which matures the 3’ end of 18S rRNA^49^ (Fig. S10b). “DNA replication” associated genes include replication factors RPA2 and putative, RPC3 (Fig S10c), both of which show growth defects in PCFs ^35,50^.

**Figure 5.**
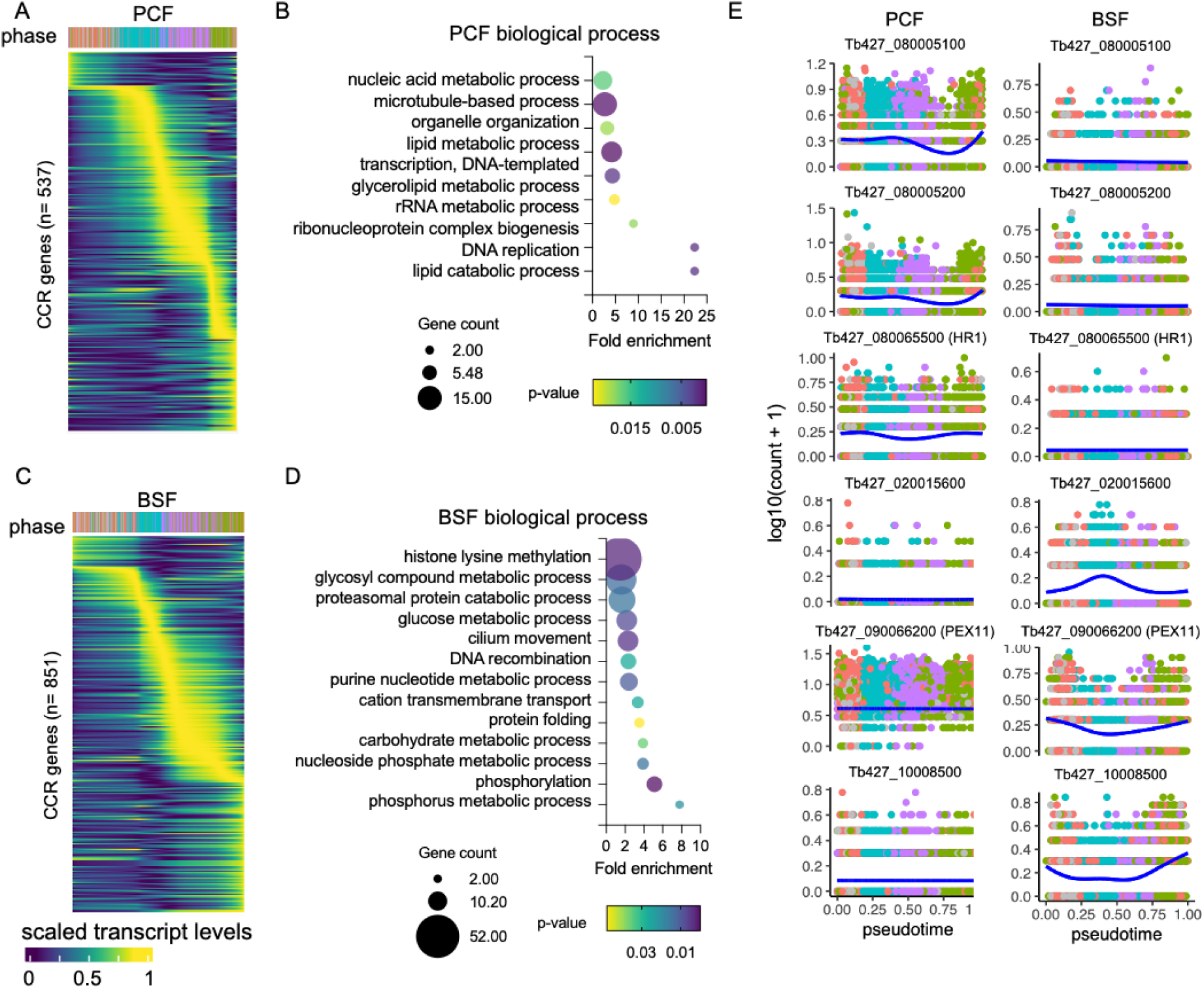
Unique CCR transcripts of BSF and PCF *T. brucei*. (**a**) Scaled transcript levels of unique CCR genes (rows), ordered by peak time, plotted across transcriptomes (columns) ordered in pseudotime across PCF cell cycle. Top annotation indicates cell phase. (**b**) GO terms associated with CCR genes unique to PCFs. Fold change of detected genes is plotted on x-axis. Points are sizes by number of genes and coloured by p-value. (**c**) Scaled transcript levels for CCR unique to BSF cells cycle, as in a. (**d**) GO terms associated with unique BSF CCR genes, as in b. (**e**) Transcript levels of 6 genes with strong association bias to one life cycle form cell cycle. Counts per cell (y-axis) are plotted across PCF (left) and BSF (right) pseudotime (x-axis), coloured by phase as in a. Blue line shows smoothed expression level across pseudotime.

The 1,294 uniquely DE genes in BSFs also showed varied expression dynamics over the cell cycle (Fig. 5c). GO term analysis highlighted 52 genes linked to the term “phosphorus metabolic process”, 12 to “carbohydrate metabolic process” and 7 specifically to “glycosyl compound metabolic process”. These metabolic associated genes include 24 components of the glycolysis/gluconeogenesis pathway, including glucose-6-phosphate isomerase (PGI), phosphoglycerate kinase (PGKC), and triosephosphate isomerase (TIM) (Fig S11a). Enzymes linked to the term “phosphorylation” included Repressor of Differentiation Kinases 1 (RDK1) and 2 (RDK2), both of which repress differentiation from BSF to PCFs^35^, and a pseudokinase linked to slowed growth in BSFs when depleted in a kinase-specific RIT-seq screen^35^. Interestingly, RDK1 and RDK2 show inverse expression patterns, with RDK1 transcripts at lower levels in late G1 before rising as the cell cycle progresses through S, G2M and back into early G1 (Fig S11b). Four genes are linked to “DNA recombination”: RPA1, KU80, RAD51 and RecQ helicase, each with a varied expression pattern (Fig S11c). Two histone-lysine n-methyltransferases, DOT1A and DOT1B, are also significantly CCR only in BSFs with these thresholds. However, both DOT1s do have a similar smoothed expression pattern in BSF and PCF (Fig. S11d) despite neither gene reaching the required thresholds in this analysis to be considered CCR in PCF (supplementary data 7).

In addition, cyclin encoding gene CYC4 was CCR only in BSFs where transcripts peak in G2/M (Fig. S9). A previously un-investigated cyclin-domain containing gene was uncovered in this analysis, Tb427_110012500, which peaks between early and late G1 phases of BSFs (Fig. S9). In contrast, CYC9 is significant only in PCFs, yet shows only a slight increase in transcript levels as the cell cycle progresses to G2/M (Fig. S9).

Thresholds for classifying CCR were selected based on the detection of previously identified cell cycle phase markers^12^ in this scRNA-seq analysis in PCFs (Fig. S4). Genes were only considered common to each form if both adjusted p-value <0.01 and FC >1.5 thresholds were satisfied in both forms, otherwise they are considered unique to one form. The transcript dynamics for six of the genes showing greatest difference in p-value between forms (Fig. S8) are plotted Fig. 5e. These include 5 genes encoding hypothetical proteins and one encoding putative peroxisomal biogenesis factor 11 (PEX11), which is only CCR in BSF and peaks in G2M and early G1 phases. If only p-values are considered when comparing BSF and PCF CCRs, 1,513 genes are considered common to both forms, 522 unique to PCFs and 366 to BSFs. If using the FC threshold only to compare genes, 1,804 are considered CCR in both, 271 in PCFs and 325 in BSFs. All comparisons are available in supplementary data 7.

## Discussion

In this work we provide the CCR transcriptomes of both BSF and PCF *T. brucei*, generated from asynchronously replicating populations. Computational reconstruction of the cell cycle with individual transcriptomes allowed us to ascertain the extent to which each gene’s transcript levels follow the periodic waves of the cell cycle and map their dynamic patterns. Comparison between transcript expression patterns and previously published protein abundance changes identified a relative delay in peak levels for transcript and protein for at least 50% of the genes that could be compared. Comparing BSF and PCF cyclic transcriptomes identified a common set of highly conserved CCR genes, enriched for known cell cycle related genes and, thus, likely novel regulators of cell cycle in kinetoplastidae. Intriguingly, a key difference between forms appears at the S-G2 transition where the gene expression switch associated with these phases is more tightly regulated in PCFs compared to BSFs.

### Cryopreservation as a method to capture transcriptomes

In addition to specific analysis of the cell cycle, we provide evidence that scRNA-seq analysis of cryopreserved parasites is feasible without detrimentally altering the transcriptome of parasites providing a methodological development likely to be of utility in multiple scRNA-seq studies.

scRNA-seq has proved a powerful method for investigating trypanosome parasites, yet its implementation is still restricted by high cost and the need to isolate live parasites^21^. We previously attempted to perform Chromium scRNA-seq with BSF *T. brucei* fixed in methanol, but this resulted in low transcripts detection per cell preventing meaningful analysis^21^. Howick *et al*. (2022) employed plate-based method SMART-seq2 to analyse *T. brucei* isolated from tsetse flies^19^. This method generally generates higher coverage transcriptomes with full-length transcripts, but at lower through-put than droplet-based methods, as this technique is limited by the use of multi-well plates to isolate cells. These authors compared the transcriptomes derived from live *T. brucei* to those prepared with two preservation methods: dithio-bis(succinimidyl propionate) (DSP) fixation or Hypothermosol-FRS preservation. Although both methods resulted in the recovery of transcriptomes comparable with live cells, conclusions could not be drawn about the impact of these methods as each was applied to samples from different experimental time points, confounding comparisons^19^.

Here, we compare Chromium generated transcriptomes from PCF and BSF prepared immediately from *in vitro* culture to those carefully cryopreserved with 10% glycerol and then thawed slowly. Comparison between fresh and frozen samples revealed few significant changes to gene expression, both when considering expression averaged across the population and between individual transcriptomes. Only one gene showed altered transcript levels between conditions for both forms: fructose-bisphosphate aldolase (ALD; Tb427_100060400), a component of the glycolytic pathway, was downregulated after freezing. ALD protein is detectable in both BSF^51^ and PCF^52,53^, but has higher transcript levels in BSFs in other studies^54–57^. It is unclear here whether the temperature changes, or use of glycerol (which BSFs have been demonstrated to use as a substrate in gluconeogenesis^58^) in the freezing/thawing procedure triggered decreased ALD transcripts. In PCFs, procyclin associated genes PAG1-5 were all upregulated in cryopreserved transcriptomes. PAGs are not essential for differentiation from BSF to PCF, but mRNA levels of PAGs 1-3 were transiently upregulated during the BSF to PCF differentiation, trigged by reducing temperature and addition of cis-aconitate^59^. PAG4 and PAG5 were not analysed in that study as levels were not detectable by blotting^59^. Hence, PAG transcript level changes are likely to be induced by the temperature change during cryopreservation.

Other than these isolated changes, we could not find significant differences between fresh and frozen transcriptomes in either form. Furthermore, freezing had little effect on the transcript recovery per cell, and samples could be fully integrated to study the biological process of interest without confounding results. Thus, cryopreservation is an appropriate method of storing *T. brucei*, and likely related parasite species, prior to scRNA-seq.

### Global cell cycle analyses

Previously, profiling of the PCF cell cycle transcriptome relied on centrifugal elutriation or serum starvation^12^. In both cases, parasites were returned to normal culture conditions and RNA was extracted from discrete time-points for sequencing. Although time points were clearly enriched for cell cycle phases, samples still contained mixed populations to varying degrees. Now technological and analytical advances have made it possible to avoid these potentially stress inducing methods by performing scRNA-seq directly on asynchronous mixed populations, with their cell cycle phases then resolved computationally. We applied pseudotime inference and differential expression methods to profile cyclical transcript changes, rather than directly comparing discretely grouped phases. Although it is likely that genes with low transcript levels are missed in this analysis, as sensitivity of scRNA-seq is lower than bulk-RNA-seq^60–62^, 1,550 genes with dynamic transcript level changes reflective of the cell cycle were identified, including 1,151 which had not been identified by bulk analysis. These CCR genes include new transcriptional markers of each phase, including those clearly distinguishing late G1 phase PCFs from early G1 phase parasites, for which previously identified early G1 markers were insufficient for labelling (Fig. 2b, f).

scRNA-seq also allowed the characterisation of the BSF cycling transcriptome for the first time. Using the same significance thresholds, we identified 1,864 genes with CCR transcript levels, 1,013 of which were also identified as CCR in the PCF cell cycle. The additional CCR genes identified only in BSF included those linked to glycolysis, which BSFs rely on to generate ATP from the glucose energy source in the mammal^63^. Interesting, the knockdown of 11 glycolysis-associated genes was linked to cell cycle arrest in G1^45^ and so further investigation may unveil if BSFs use glycolysis activity levels as a signal for re-entering the cell cycle during G1. Other genes uniquely CCR in BSFs include DNA recombination factors, RecQ helicase. RECQ functions to repair DNA breaks, including at the subtelomeric sites of variant surface glycoprotein (VSG) expression^64^, and is hypothesised to limit strand exchange during homologous recombination (HR) reactions at this site^65^. HR at *VSG* expression sites is central to antigenic variation required for evasion of the mammalian immune system and so survival of BSF parasites. RAD51 is central to the recombination of previously silent *VSGs* into the transcribed *VSG* expression site to allow expression of a new VSG on the parasite surface^66^. It is hence possible that these genes show higher CCR expression in BSFs due to their role in antigenic variation-associated HR events, which may be triggered by DNA replication-associated damage^67^ and so require specific expression timing in the cell cycle.

A further notable difference between forms is the clear distinction of S and G2M phases in PCFs, compared to much less apparent separation in BSFs when using both UMAP and independent pseudotime inference approaches. This indicates that the switch in gene expression associated with the S-G2 transition is much more discrete or tightly regulated in PCFs than BSFs. Comparing the expression patterns of shared CCR genes in each form (Fig. 4) further highlights that expression patterns of the G1 and S phase genes are highly comparable between forms, whereas after S phase the timing of gene expression is far less synchronised. Human cells display tight regulation of the S-G2 transition, with the mitotic gene network only expressing after the end of S phase^68^. Here, ATR kinase remains in its active form throughout S phase and cells only progress to G2, and upregulate the associated gene programme, upon ATR inactivation at the end of S to ensure complete genome replication prior to mitosis^68^. In the absence of ATR, human cells activate DNA replication origin firing aberrantly, and undergo premature and defective mitosis^69^.

Interestingly, ATR activity in *T. brucei* is required for normal S phase progression in both PCFs^70^ and BSFs^71^, yet the proteins’ role in the S-G2 transition differs dramatically between forms. In BSFs, ATR depletion is lethal and within 24 hours increases the proportions of S and G2M phase parasites, as well as aberrant cells resulting from premature mitosis and cytokinesis events, indicating a putatively similar role to human ATR^71^. Yet in PCFs, ATR knockdown has little effect on the cell cycle indicating PCFs mostly undergo the S-G2 transition and complete mitosis and cytokinesis correctly without ATR activity^70^. Thus, as highlighted by scRNA-seq investigations here, PCFs and BSFs appear to, at least partially, regulate the S-G2 transition differently. Why BSFs would not require the same level, or mechanism, of regulation of this transition is currently unclear. Intriguingly however, even in the presence of persistent DNA damage BSFs will continue to replicate DNA and proliferate^72^. As BSFs require DNA damage at VSG expression sites to trigger HR and VSG switching, it is plausible that BSF allow continuation to G2 in the presence of DNA damage acquired during S phase, which could then be repaired to facilitate VSG recombination event in subsequent phases. Indeed, Rad51 transcripts peak at the S-G2M transition, a pattern not observed in PCFs (Fig. S11c), and in BSFs Rad51 foci form mainly in G2/M phase parasites^73^.

### Conservation of cell cycles between life stages

Interrogating the shared and unique CCR transcriptomes is likely to unveil new insights into *T. brucei* cell cycle regulation, for example by assess expression patterns of cyclins. In *T. brucei*, 13 cyclins have been investigated and several cyclin-CRK binding pairs have been documented^6,11,74,75^. Notably, we find just two cyclins with strong CCR transcript dynamics in both forms, CYC8 and CYC6. CYC6 binds CRK3^76^ and is well characterised as essential for mitosis^76–78^ in both forms, correlating with expression levels detected here at the S-G2M transition. CYC8 instead clearly peaks during late G1, despite RNAi depletion leading to a slight increase in G2/M cells in PCFs^77^. Thus, although both cyclins have roles in G2/M, CYC8 peak earlier in the cell cycle and is followed by the gradual rise in CYC6. Despite these different patterns, transcripts of both cyclins are reported to be bound by the RNA binding protein RBP10^79^, highlighting that steady state RNA levels are likely regulated by multiple factors beyond the individual RBPs. Additionally, RBP10 is not expressed in PCF^80,81^, and so how matching cyclic expression patterns are regulated in both forms is unclear. CYC8 transcripts were previously identified as enriched in G1^12^, yet protein levels were undetectable^13^. Protein levels of CYC6 have been documented as CCR^13^, yet previously, CYC6 transcripts were not recorded as CCR^12^, exemplifying the power of scRNA-seq over bulk transcriptomics. ScRNA-seq analysis finds only a slight CYC9 transcript increase in G2M, and only in PCFs. Yet, in BSFs CYC9 transcript depletion results in a clear cytokinesis defeat^82^. Thus, transcript FC does not necessarily correlate with functional significance, as was also noted when comparing CCR genes to cell cycle defects profiled by the genome-scale screen in BSFs^45^. Results of CYC9 RNAi depletion in PCFs is currently conflicting, possibly due to differences in knockdown efficiency, as one study observed a substantial cell cycle arrest in G2/M^77^, while another saw no specific arrest in any cell cycle phase^82^. In both forms CYC4 transcript levels dip in late G1 before rising again in S phase through to G2M, but only reached a FC > 1.5 in BSFs. Interestingly, RNAi against CYC4 in PCF highlighted the cyclin’s role in the G1/S transition^83^, again indicating transcript regulation does not predict phenotypic outcome. Finally, a novel putative cyclin, Tb427_110012500, was detected with CCR transcripts in BSF form only, where transcripts peak between early and late G1. This gene contains a cyclin N-terminal domain, but no functional analysis has been published. Of the remain 9 documented cyclins^6,11,84^ in *T. brucei*, none reached significance thresholds in either form.

### Transcript and protein periodicity

Lastly, we compared transcript and protein abundance levels across the cell cycle. In the human cell cycle, just 15% of CCR proteins are encoded by genes which also have CCR transcripts^85^. In this study we also observed little correlation between transcript and protein regulation during the *T. brucei* cell cycle. Thus, for most genes the cyclic protein abundance patterns are the result of mostly translational, and post-translation processes. Even accounting for experimental differences in approaches, why so many transcripts show cyclic expression patterns without resulting in significant protein changes, especially in the absence of transcriptional control due to polycistronic transcription in *T. brucei*^86,87^, remains a puzzling question across eukaryotes. Of those genes that were identified as CCR for both transcript and protein abundance, a relative delay was observed for the majority of genes. A time delay between peak transcript and proteins levels was also observed in human cells^85^. Such a delay may allow *T. brucei* to prepare for the subsequent phase by upregulating transcripts, after which translation can rapidly generate the required proteins. A similar observation can be made during *T. brucei* life cycle progression: stumpy bloodstream forms upregulate hundreds of transcripts related to PCF biology^20,55,56,88,89^ in preparation for differentiation, but not all upregulated genes are detectable in proteomic analysis of stumpy forms and instead appear after the rapid development of PCFs once the environmental trigger to differentiate has been received^80,90^.

In summary, the experiments discussed here exploit cryopreservation to preserve *T. brucei* for scRNA-seq analysis, an approach that can be likely also be extended to and related parasites, to increase flexibility and feasibility of experimental design. Making use of these data we have generate detailed transcriptome atlases of the BSF and PCF cell cycles, which can be further interrogated by the publicly accessible interactive webtool (https://cellatlas-cxg.mvls.gla.ac.uk/Tbrucei.cellcycle.bsf/ and https://cellatlas-cxg.mvls.gla.ac.uk/Tbrucei.cellcycle.pcf/).

## Materials and Methods

### *T. brucei* culture

For scRNA-seq experiments bloodstream form Lister 427 were cultured in HMI-9^91^ with 20% foetal calf serum (FCS) at 37 °C with 5% CO_2_. Procyclic form Lister 427 were cultured in SDM-79^92^ supplemented with 10% FCS and 0.2% hemin, at 27 °C in sealed flasks without CO_2_. A haemocytometer was used for all cell density and motility counts. For mNeonGreen tagging experiments Lister 427 BSF expressing Cas9 were used (gifted, R. McCulloch), these were transfected with J1339 plasmid^93^, which allows constitutive expression of Cas9.

### scRNA-seq sample preparation with cryopreserved *T. brucei*

For cryopreservation of both PCF and BSF, fresh 2X freezing media with FCS and 20% glycerol was used for each sample. Cell density was adjusted to 2×10^6^/ml before parasite culture and 2X freezing media were mixed 1:1 by slow addition of freezing media to culture and gentle resuspension. Cells were aliquoted into 1ml cryopreservation tubes, wrapped in cotton wall to prevent rapid cooling and incubated at −80 °C for 24 hr. Tubes were then moved to LN_2_ storage. 1 ml samples were thawed immediately before scRNA-seq library preparation. Tubes were placed at room temperature (RT) for 5-10 min before incubating at 37 °C (BSF) or 27°C (PCF), until a small ice crystal was left in the tube. Cells were moved to RT until completely defrosted then pipetted with wide-bore pipette tips into 50 ml falcons. 1 ml of pre-warmed media with FCS (37 °C or 27°C as appropriate) was added drop-wise to falcon with and swirled gently. A further 1ml of media was used to rinse the cryotube with a wide-bore tip and added drop-wise to falcon. Increasing volumes of pre-warmed media was added to the cells (3 ml, 6ml and 12 ml) drop-wise, with at least 1 min pause between each addition. Cells were pelleted by centrifugation at 400 x g, for 10 min at RT, and the supernatant was poured off. 10 ml of media was then added dropwise to wash cells. Cells were centrifuged again and supernatant poured off, before resuspending in 1 ml of 1X PBS supplemented with 1% D-glucose (PSG) and 0.04% bovine serum albumin (BSA), by gentle pipetting. Cells were strained through a 40 μm filter into a 1.5 ml Eppendorf. Cells were centrifuged at 400 x g for 10 min at RT and supernatant was removed with a pipette. Cells were suspended in 150 µl of PSG + 0.04% BSA. Sample was diluted 1:1 and in PSG + 0.04% BSA and loaded to haemocytometer to determine cell concentration. Cell concentration was adjusted to 1,000 cells/µl and stored on ice.

### scRNA-seq sample preparation of fresh *in vitro* cultured *T. brucei*

For both BSF and PCF *T. brucei*, 1×10^6^ cells were transferred to a falcon tube and were centrifuged at 400 x g for 10 min at RT. The supernatant was poured off and pre-warmed media added dropwise to the sample to was cells. Cells were centrifuged again and supernatant poured off, before resuspending in 1 ml of PSG + 0.04% BSA, by gentle pipetting with wide-bore pipette tips. Cells were strained through a 40 μm filter into a 1.5 ml Eppendorf before centrifuging again at 400 x g for 10 min at RT removing the supernatant with a pipette. Cells were suspended in 150 µl of PSG + 0.04% BSA and concentration adjusted to 1,000 cells/ul before storing on ice.

### Chromium (10x Genomics) library preparation and Illumina sequencing

As BSF and PCF as easily identified by known transcriptional differences^54–57,89,94^, the two forms were multiplexed. Fresh BSF and PCF were combined in approximate equal ratio into sample 1, and cryopreserved BSF and PCF into sample 2. 14,000 cells of each sample were loaded onto the Chromium Control and library preparation was performed with the Chromium Single Cell 3’ chemistry version 3.1 kits. (*L. major* parasites were additionally multiplexed with each sample as performed previously with kinetoplastids^20^, but are not analysed here). Libraries were sequences with Illumina NextSeq 2000, to generate 28 bp x 130 bp paired reads to a depth of 46,561 and 43,332 mean reads per cell for sample 1 and sample 2, respectively. Library preparation and sequencing was performed by Glasgow Polyomics.

### Data mapping and count matrix generation

To improve the proportion of mapped reads attributed to a feature for transcript counting, the UTR annotation of the Lister 427 2018 reference genome^47^ were extended. 2,500 bp were added to the end each annotated coding region of the gtf file (unless the annotation overlapped with the next genomic feature in which case the UTR was extended to the base before the next feature). The same approach was used to edit the *L. major* Friedlin reference genome annotation^95^. Reads were mapped to both the *T. brucei* WT427 2018 and *L. major* Friedlin references and counts matrix generated with Cell ranger v 7. *L. major* transcriptomes and those of multiplets containing transcripts from both species were removed from analysis. The resulting count matrices and samples summaries are available in supplementary data 1 and at Zenodo (10.5281/zenodo.7508131).

### Sample de-multiplexing and QC filtering

To de-multiplex the PCF and BSF transcriptomes, a set of high confidence marker genes was defined from published bulk-RNA-seq studies where two replicates are available for DE analysis. DE analysis was performed using TriTrypDB^96^ which implements DESeq2^97^ to compare datasets. DE between Lister 427 PCFs and Lister427 monomorphic BSFs^57^, and slender pleomorphic BSF EATRO 1125 (clone AnTat 1.1) and experimentally derived early PCFs^89^, identified 238 BSF and 221 PCF marker genes (FC > 2, p-value < 0.05, supplementary data 1). As PCF and BSFs were expected to be present at around a 1:1 ratio, marker genes detected in 20% - 70% of the cells were selected as markers (Supplementary data 1). This gave 157 and 50 high confidence marker genes for PCFs and BSFs, respectively. scGate^98^ was used to gate BSF, PCF, multiplets containing a mix of each life cycle form using marker genes and transcriptomes not enriched for either form (supplementary data 1). Once, demultiplexed into each sample (BSF fresh, BSF frozen, PCF fresh and PCF frozen) cells were filtered for homogenous multiples with higher-than-average UMI and feature counts, and poor-quality transcriptomes with low UMI and feature counts (Fig. 1a, b). Finally, cells expressing higher than average mitochondrial transcripts encoded on the kDNA maxi circle were removed, as likely generated with lysing cells (Fig. 1c). Full code is available for all steps at Zenodo (10.5281/zenodo.7508131)

### Live vs cryopreserved *T. brucei* differential expression analysis

The AverageExpression function from Seruat v4.1.0^99^ was used to average expression of each gene across cells for each sample. Fold changed was calculated as average expression for frozen over fresh samples for each life cycle form separately. Genes with average expression < 0.05 in each fresh or frozen were excluded from fold change analysis. For PCA analysis, data was “pseudobulked” by summing counts across all cells for each gene, per condition. DESeq2 v1.32.0 was used to log2 scale counts and generate PCA plot. DE analysis between individual transcriptomes of each condition was performed with Seurat function FindAllMarkers using MAST test^100^. Only genes detected in 25% of cells in tested condition and with FC > 1.5 between fresh and frozen were considered.

### Data integration and dimensional reduction

Each sample was normalised and log2 transformed using Scran v1.22.1, as described previously^20^. The top 3,000 variable genes were identified in each sample using two independent methods (Scran, which using log2 counts and Seurat applied to raw counts), and results compared to select common variable genes. 1,939 genes were identified for BSF fresh, 2,063 for BSF frozen 2,000 for PCF fresh and 1,924 for PCF frozen (supplementary data 1). For data integration variable genes for fresh and frozen samples were compared and selected using SelectIntegrationFeatures, and filtered for only those with standardised variance over 1 in both conditions. BSF and PCF samples were considered separately, identifying 1,652 and 1,738 variable genes for integration, respectively (supplementary data 1). Integration was performed with Fast mutual nearest neighbours (FastMNN), which performs batch correction by finding MNN pairs of cells between conditions with mutually similar gene expression and calculating correction between these pairs. MNN does not assume equal population composition between sample and only performs correction between the overlapping subsets of cells^101^. FastMNN first performs a PCA across all cells and finds MNN between cells in this deduced dimensional space to increase speed and remove noise. The default of 50 dimensions was used to integrate fresh and frozen samples for BSF and PCF independently, and nearest-neighbours were identified for 5% of cells in each case. Integrated cells were visualised using UMAP^25^ applied to the first 30 dimensions calculated by FastMNN, implemented by the Seurat package.

### Cell cycle phase labelling

Cell cycle phases were inferred using marker gene identified with bulk RNA-seq previously^12^. Syntenic orthologs for each phase marker (originally identified in the TRUE927 genome) were found of Lister 427 2018 reference genome via TritrypDB^96^, and those detected in at least 10% of transcriptomes were selected for PCF and BSF integrated datasets independently. An ‘expression score’ for each phase was found of each using MetaFeature function from Seurat using markers. The ratio of a cell’s expression score over the mean expression scores across cells was calculated for each phase. The phase with the highest ratio was assigned to each cell. If a cell has an expression score <1 for all phase (i.e. no enrichment over the average phase score), the cell was assigned “unlabelled”.

### Pseudotime inference and differential expression analysis

For pseudotime inference the autoencoder approach from Cyclum^26^ was used for BSF and PCF separately. Raw counts for the same variable genes used for integration steps, described above, were first scaled before the model was trained using 25% of total cells and default parameters. The model was then applied to the whole dataset to infer pseudotime values for each cell. To allow clear visualisation and comparison between PCF and BSF cell cycles, pseudotime was scaled between 0 and 1 for each form and in the case of PCFs, pseudotime was shifted to set 0 to be at approximated early G1.

PseudotimeDE^27^ was used for DE analysis over pseudotime. This package calculates accurate p-values by accounting for uncertainty in the pseudotime inference, and shows greater power and lower false discovery rates (FDR) than similar packages^27^. To calculate pseudotime uncertainty, the same Cyclum training model was applied to 100 subsets of the data, each containing 80% of the cells selected at random. Genes that were detected in at least 10% of the cells were assessed for DE over pseudotime using a negative binomial generalized additive model (NB-GAM) and default settings. The empirical p-values calculated by PseudotimeDE (which take into account pseudotime uncertainty) was adjusted using the Benjamini and Hochberg method to find the FDR^102^. PseudotimeDE was applied to the normalised log-transformed counts, to account for overall increase in RNA across the cell cycle (Fig. S3).

For calculating FC in gene expression over pseudotime, the smoothed expression of each gene was predicted from the GAM fitted by PseudotimeDE. The ratio of the maximum value in this prediction over the minimum value was calculated as the FC in the average expression over pseudotime. Genes were considered CCR if adjusted p-value was below 0.01, and FC was over 1.5, based on the detection of known CCR genes^12^ (Fig. S4). Predicted models were also used when plotting smoothed expression. All GO term enrichment analysis was performed using the TriTrypDB resource^96^.

### Dataset comparison

For comparison with proteomics^13,14^, bulk transcriptomics^12^ and genome scale cell cycle defect RNAi screen^45^, all of which used the TRUE927 reference genome^103^, syntenic orthologs were identified in the Lister 427 2018 reference^47^ using TriTrypDB implementation of OrthoMCL^104^. For each study, CCR genes were retained as those selected by original authors.

### Gene conservation analysis

Orthogroups were identified for each CCR gene, common to both BSF and PCF cell cycles, and orthologous protein sequences across 44 kinetoplastida proteomes were extracted from previous analysis^46^. A distance matrix was created from orthologous protein sequences with ClutalOmega^105^. Using the distance matrix, FastME^106^ was used to calculate the tree length for each orthogroup which contained four or more protein sequences.

### Expression profiling of mNeonGreen tagged proteins

CRISPR/Cas9 editing was used to added epitope tags to three genes in BSF WT427/Ca9 (Tb427_080028700, Tb427_060036900 and Tb427_110169500). Gene specific primers were used to amplify the donor fragment containing mNeonGreen and G418 resistance gene from a pPOTv7 plasmid as previously designed^28,107^. Primers were designed using the LeishGEdit.net resource^28,107^. 30 ng circular plasmid, 0.2 mM dNTPs, 2 µM each of gene-specific forward and reverse primers and 1 unit Phusion polymerase (NEB) were mixed in 1X HF Phusion buffer and 3% (v/v) DMSO, up 50 µl total volume with H_2_0. The PCR was run as follows: 5 min at 98 °C, 40 cycles of 98 °C for 30 seconds, 65 for °C, and 72 °C for 2 min 15 seconds, followed by a final extension at 72 °C for 7 min. To amplify the sgRNA, 2 µM of gene specific forward primer, 2 µM of the generic G00 primer^28^, 0.2 µM of dNTPs, 1 unit of Phusion Polymerase were mixed with 1x HF Phusion buffer (NEB) and made up to 50 µl total volume with H_2_0. The PCR was run as follows: 98 °C for 30 seconds, followed 35 cycles of 98 °C 10 seconds, 60 °C for 30 seconds and 72 °C for 15 seconds. 2 µl of each product was run on 1% agarose gel to confirm expected size and the products were both ethanol precipitated and eluted into 5 µl of H_2_O. 1×10^7^ WT427/Cas9 BSFs were transfected in 100 µl of transfection buffer (90mM NaH_2_PO4, 5mM KCl, 150 µM CaCl_2_ and 500 mM HEPES, pH7.3) plus the 5 µl donor and 5 µl sgRNA, using the Nucleofector™ 2b Device (Lonaz) using program X-100. Parasites were serially diluted and aliquoted into 24 well plated. G418 selection was added after 16-24 hr at final concentration of 2 µg/ml and clones were recovered after 5-7 days. To confirm tag integration with PCR, genomic DNA was extracted from WT427 cas9 BSFs and three clonal derivatives for each gene using the DNeasy Blood and Tissue extraction kit (Qiagen). 5 µM each of gene-specific forward and reverse primers and 30 ng of gDNA was mixed with 0.4 µl Phire Hot Start II polymerase, 1X Phire Reaction Buffer, 0.2 mM dNTPs and up to 20 µl H_2_O. The 2-step PCR was run as follows: 30 seconds at 98 °C, 30 cycles of 98 °C for 5 seconds and 72 °C for 1 min, followed by a final extension at 72 °C for 1 min. For fluorescence and flow cytometry assays, cells were harvested by centrifugation at 400 x g for 10 mins, washed in 1 X Phosphate Buffered Saline (PBS) and fixed in 1% formaldehyde for 10 min at room temperature. Cells were pelleted and washed again in 1X PBS to remove formaldehyde. For microscopy, cells were attached to a poly-L-lysine treated slide before 5 μl of Fluoromount G with DAPI (Cambridge Bioscience, Southern Biotech) was added and coverslip applied. For flow cytometry, formaldehyde fixed cells were resuspended in 1X PBS supplemented with 5 mM EDTA and 0.1 ug/ml DAPI and incubated on ice for 30 min. DAPI and mNeonGreen fluorescence was detected for 10,000 events per sample.

## Data availability

The transcriptome data generated in this study have been deposited in the European Nucleotide Archive with project accession number PRJEB58781. The processed transcript count data and cell metadata generated in this study are available at Zenodo (10.5281/zenodo.7508131). BSF and PCF cell cycle transcriptomes can also explored using the interactive cell atlas (https://cellatlas-cxg.mvls.gla.ac.uk/Tbrucei.cellcycle.bsf/ and https://cellatlas-cxg.mvls.gla.ac.uk/Tbrucei.cellcycle.pcf/).

## Code availability

All code and necessary intermediate files are available at Zenodo (10.5281/zenodo.7508131)

## Supplementary data

Supplementary figures.pdf

Supplementary data 1 – sample processing

Tab 1. – Genes used for BSF and PCF gating

Tab 2. – Results of BSF and PCF gating

Tab 3. – scRNA-seq sample summary

Tab 4. – variable genes per sample

Supplementary data 2 – fresh vs frozen DE analysis

Tab 1. – BSF average expression comparison

Tab 2. – PCF average expression comparison

Tab 3. – BSF MAST DE test

Tab 4. – PCF MAST DE test

Supplementary data 3 – Early G1 vs unlabelled cells comparison

Tab 1. – PCF DE

Tab 2. – BSF DE

Supplementary data 4 – PCF cell cycle DE analysis

Tab 1. – PseudotimeDE analysis

Tab 2. – PseudotimeDE results for CCR genes only

Tab 3. – Average expression of each CCR per phase

Tab 4. – Dataset comparison of CCR genes

Tab 5. – Comparison with Crozier proteomics study

Tab 6. – Comparison with Benz proteomics study

Supplementary data 5 – BSF cell cycle DE analysis

Tab 1. – PseudotimeDE analysis

Tab 2. – PseudotimeDE results for CCR genes only

Tab 3. – Average expression of each CCR per phase

Tab 4. – Comparison with Crozier proteomics study

Tab 5. – Comparison with Benz proteomics study

Supplementary data 6 – Common CCR gene analysis

Tab 1. – CCR comparison between BSF and PCF

Tab 2. – Comparison of peak expression between BSF and PCF

Tab 3. – Ortholog groups and conservation of common CCR genes

Tab 4. – Comparison of CCR genes and cell cycle defects

Tab 5. – Early G1 phase associated GO terms

Tab 6. – Late G1 phase associated GO terms

Tab 7. – S phase associated GO terms

Tab 8. – G2M phase associated GO terms

Tab 9. – Plotted GO terms

Supplementary data 7 – Unique CCR gene analysis

Tab 1. – GO terms associated with CCR unique to PCF

Tab 2. – GO terms associated with CCR unique to BSF

## Acknowledgements

We thank J. Galbraith and P. Herzyk (Glasgow Polyomics, University of Glasgow) for their guidance, library preparation, sequencing and data handling. We also thank R. McCulloch (University of Glasgow) for provision of cell lines and manuscript comments, E. Agboraw (University of Glasgow) for uploading data to the webtool. This research was funded in part by the Wellcome Trust [Grant numbers 218648/Z/19/Z awarded to E.M.B., 104111/Z/14/ZR to T.D.O., 221717/Z/20/Z to K.R.M. and 220058/Z/19/Z awarded to K.R.M in support of G.R.O] and the Biotechnology and Biological Sciences Research Council [Grant numbers BB/R017166/1 awarded to R. McCulloch in support of C.A.M and BB/W001101/1 awarded to R. McCulloch and C.A.M]. For the purpose of open access, the author has applied a CC BY public copyright licence to any Author Accepted Manuscript version arising from this submission.

## Author contributions

Methodology: E.M.B., C.A.M., G.R.O., J.H, T.D.O, and K.R.M. Data collection: E.M.B and C.M. Bioinformatic data analysis: E.M.B and G.R.O. All authors participated in discussions related to this work. All authors wrote, reviewed and approved the manuscript.

## References

1. Matthews, K. R. The developmental cell biology of Trypanosoma brucei. J. Cell Sci. 118, 283–90 (2005).

2. Christiano, R. et al. The proteome and transcriptome of the infectious metacyclic form of Trypanosoma brucei define quiescent cells primed for mammalian invasion. Mol. Microbiol. 106, 74–92 (2017).

3. Rojas, F. & Matthews, K. R. Quorum sensing in African trypanosomes. Curr. Opin. Microbiol. 52, 124–129 (2019).

4. Matthews, K. R. Trypanosome Signaling-Quorum Sensing. Annu. Rev. Microbiol. 75, 495–514 (2021).

5. Silvester, E., McWilliam, K. R. & Matthews, K. R. The cytological events and molecular control of life cycle development of Trypanosoma brucei in the mammalian bloodstream. Pathogens vol. 6 (2017).

6. Hammarton, T. C. Cell cycle regulation in Trypanosoma brucei. Mol. Biochem. Parasitol. 153, 1–8 (2007).

7. Li, Z. Regulation of the cell division cycle in Trypanosoma brucei. Eukaryot. Cell 11, 1180–1190 (2012).

8. Wheeler, R. J., Gull, K. & Sunter, J. D. Coordination of the Cell Cycle in Trypanosomes. https://doi.org/10.1146/annurev-micro-020518-115617 73, 133–154 (2019).

9. Passos, A. de O. et al. The Trypanosomatids Cell Cycle: A Brief Report. 25–34 (2022) doi:10.1007/978-1-0716-2736-5_2/FIGURES/1.

10. Mottram, J. C. & Smith, G. A family of trypanosome cdc2-related protein kinases. Gene 162, 147–152 (1995).

11. Lee, K. J. & Li, Z. The CRK2-CYC13 complex functions as an S-phase cyclin-dependent kinase to promote DNA replication in Trypanosoma brucei. BMC Biol. 19, 1–15 (2021).

12. Archer, S. K., Inchaustegui, D., Queiroz, R. & Clayton, C. The Cell Cycle Regulated Transcriptome of Trypanosoma brucei. PLoS One 6, e18425 (2011).

13. Crozier, T. W. M. et al. Proteomic Analysis of the Cell Cycle of Procylic Form Trypanosoma brucei. Mol. Cell. Proteomics 17, 1184–1195 (2018).

14. Benz, C. & Urbaniak, M. D. Organising the cell cycle in the absence of transcriptional control: Dynamic phosphorylation co-ordinates the Trypanosoma brucei cell cycle post-transcriptionally. PLoS Pathog. 15, (2019).

15. Tritschler, S. et al. Concepts and limitations for learning developmental trajectories from single cell genomics. Development 146, (2019).

16. Wolfien, M., David, R. & Galow, A.-M. Single-Cell RNA Sequencing Procedures and Data Analysis. Bioinformatics 19–35 (2021) doi:10.36255/EXONPUBLICATIONS.BIOINFORMATICS.2021.CH2.

17. Hutchinson, S. et al. The establishment of variant surface glycoprotein monoallelic expression revealed by single-cell RNA-seq of Trypanosoma brucei in the tsetse fly salivary glands. PLoS Pathog. (2021) doi:10.1371/journal.ppat.1009904.

18. Vigneron, A. et al. Single-cell RNA sequencing of Trypanosoma brucei from tsetse salivary glands unveils metacyclogenesis and identifies potential transmission blocking antigens. Proc. Natl. Acad. Sci. U. S. A. 117, 2613–2621 (2020).

19. Howick, V. M. et al. Single-cell transcriptomics reveals expression profiles of Trypanosoma brucei sexual stages. PLOS Pathog. 18, e1010346 (2022).

20. Briggs, E. M., Rojas, F., McCulloch, R., Matthews, K. R. & Otto, T. D. Single-cell transcriptomic analysis of bloodstream Trypanosoma brucei reconstructs cell cycle progression and developmental quorum sensing. Nat. Commun. (2021) doi:10.1038/s41467-021-25607-2.

21. Briggs, E. M., Warren, F. S. L., Matthews, K. R., Mcculloch, R. & Otto, T. D. Application of single-cell transcriptomics to kinetoplastid research. Parasitology (2021) doi:10.1017/S003118202100041X.

22. Smith, T. K., Bringaud, F., Nolan, D. P. & Figueiredo, L. M. Metabolic reprogramming during the Trypanosoma brucei life cycle. F1000Research 6, (2017).

23. Benne, R. Mitochondrial genes in trypanosomes. Trends Genet. 1, 117–121 (1985).

24. Finak, G. et al. MAST: a flexible statistical framework for assessing transcriptional changes and characterizing heterogeneity in single-cell RNA sequencing data. Genome Biol. 16, 278 (2015).

25. McInnes, L., Healy, J. & Melville, J. UMAP: Uniform Manifold Approximation and Projection for Dimension Reduction. (2018).

26. Liang, S., Wang, F., Han, J. & Chen, K. Latent periodic process inference from single-cell RNA-seq data. Nat. Commun. 2020 111 11, 1–8 (2020).

27. Song, D. & Li, J. J. PseudotimeDE: inference of differential gene expression along cell pseudotime with well-calibrated p-values from single-cell RNA sequencing data. Genome Biol. 22, 1–25 (2021).

28. Beneke, T. et al. A CRISPR Cas9 high-throughput genome editing toolkit for kinetoplastids. R. Soc. Open Sci. 4, 170095 (2017).

29. Esson, H. J. et al. Morphology of the trypanosome bilobe, a novel cytoskeletal structure. Eukaryot. Cell 11, 761–772 (2012).

30. Morriswood, B. & Schmidt, K. A MORN Repeat Protein Facilitates Protein Entry into the Flagellar Pocket of Trypanosoma brucei. Eukaryot. Cell 14, 1081–1093 (2015).

31. Hines, J. C. & Ray, D. S. A mitochondrial DNA primase is essential for cell growth and kinetoplast DNA replication in Trypanosoma brucei. Mol. Cell. Biol. 30, 1319–1328 (2010).

32. Barandun, J. et al. The complete structure of the small-subunit processome. Nat. Struct. Mol. Biol. 2017 2411 24, 944–953 (2017).

33. Tu, X. & Wang, C. C. Pairwise Knockdowns of cdc2-Related Kinases (CRKs) in Trypanosoma brucei Identified the CRKs for G1/S and G2/M Transitions and Demonstrated Distinctive Cytokinetic Regulations between Two Developmental Stages of the Organism. Eukaryot. Cell 4, 755 (2005).

34. Tu, X. & Wang, C. C. The Involvement of Two cdc2-related Kinases (CRKs) in Trypanosoma brucei Cell Cycle Regulation and the Distinctive Stage-specific Phenotypes Caused by CRK3 Depletion. J. Biol. Chem. 279, 20519–20528 (2004).

35. Jones, N. G. et al. Regulators of Trypanosoma brucei cell cycle progression and differentiation identified using a kinome-wide RNAi screen. PLoS Pathog. 10, e1003886 (2014).

36. Tu, X., Kumar, P., Li, Z. & Wang, C. C. An aurora kinase homologue is involved in regulating both mitosis and cytokinesis in Trypanosoma brucei. J. Biol. Chem. 281, 9677–9687 (2006).

37. Boynak, N. Y. et al. Identification of a Wee1–Like Kinase Gene Essential for Procyclic Trypanosoma brucei Survival. PLoS One 8, e79364 (2013).

38. Dang, H. Q. & Li, Z. The Cdc45·Mcm2–7·GINS Protein Complex in Trypanosomes Regulates DNA Replication and Interacts with Two Orc1-like Proteins in the Origin Recognition Complex. J. Biol. Chem. 286, 32424–32435 (2011).

39. Li, Z. & Wang, C. C. KMP-11, a Basal Body and Flagellar Protein, Is Required for Cell Division in Trypanosoma brucei. Eukaryot. Cell 7, 1941 (2008).

40. Ishii, M. & Akiyoshi, B. Characterization of unconventional kinetochore kinases KKT10 and KKT19 in Trypanosoma brucei. J. Cell Sci. 133, (2020).

41. Zhou, Q. et al. Faithful chromosome segregation in Trypanosoma brucei requires a cohort of divergent spindle-associated proteins with distinct functions. Nucleic Acids Res. 46, 8216 (2018).

42. Zhou, Q., Hu, H. & Li, Z. KLIF-associated cytoskeletal proteins in Trypanosoma brucei regulate cytokinesis by promoting cleavage furrow positioning and ingression. J. Biol. Chem. 298, 101943 (2022).

43. Bessat, M., Knudsen, G., Burlingame, A. L. & Wang, C. C. A Minimal Anaphase Promoting Complex/Cyclosome (APC/C) in Trypanosoma brucei. PLoS One 8, (2013).

44. Manzano-López, J. & Monje-Casas, F. The Multiple Roles of the Cdc14 Phosphatase in Cell Cycle Control. Int. J. Mol. Sci. 2020, Vol. 21, Page 709 21, 709 (2020).

45. Marques, C. A., Ridgway, M., Tinti, M., Cassidy, A. & Horn, D. Genome-scale RNA interference profiling of Trypanosoma brucei cell cycle progression defects. Nat. Commun. 2022 131 13, 1–16 (2022).

46. Oldrieve, G. R., Malacart, B., López-Vidal, J. & Matthews, K. R. The genomic basis of host and vector specificity in non-pathogenic trypanosomatids. Biol. Open 11, (2022).

47. Müller, L. S. M. et al. Genome organization and DNA accessibility control antigenic variation in trypanosomes. Nature (2018) doi:10.1038/s41586-018-0619-8.

48. Jaremko, D., Ciganda, M. & Williams, N. Trypanosoma brucei Homologue of Regulator of Ribosome Synthesis 1 (Rrs1) Has Direct Interactions with Essential Trypanosome-Specific Proteins. mSphere 4, (2019).

49. Kala, S. et al. The interaction of a Trypanosoma brucei KH-domain protein with a ribonuclease is implicated in ribosome processing. Mol. Biochem. Parasitol. 211, 94–103 (2017).

50. Rocha-Granados, M. C. et al. Identification of proteins involved in Trypanosoma brucei DNA replication fork dynamics using nascent DNA proteomics. bioRxiv 468660 (2018) doi:10.1101/468660.

51. Leite, A. B. et al. Effect of lysine acetylation on the regulation of Trypanosoma brucei glycosomal aldolase activity. Biochem. J. 477, 1733–1744 (2020).

52. Jones, A. et al. Visualisation and analysis of proteomic data from the procyclic form of Trypanosoma brucei. Proteomics 6, 259–267 (2006).

53. Vertommen, D. et al. Differential expression of glycosomal and mitochondrial proteins in the two major life-cycle stages of Trypanosoma brucei. 158, 189–201 (2008).

54. Siegel, T. N., Hekstra, D. R., Wang, X., Dewell, S. & Cross, G. A. M. Genome-wide analysis of mRNA abundance in two life-cycle stages of Trypanosoma brucei and identification of splicing and polyadenylation sites. Nucleic Acids Res. 38, 4946–4957 (2010).

55. Kabani, S. et al. Genome-wide expression profiling of in vivo-derived bloodstream parasite stages and dynamic analysis of mRNA alterations during synchronous differentiation in Trypanosoma brucei. BMC Genomics 10, 427 (2009).

56. Queiroz, R., Benz, C., Fellenberg, K., Hoheisel, J. D. & Clayton, C. Transcriptome analysis of differentiating trypanosomes reveals the existence of multiple post-transcriptional regulons. BMC Genomics 10, 495 (2009).

57. Jensen, B. C. et al. Extensive stage-regulation of translation revealed by ribosome profiling of Trypanosoma brucei. BMC Genomics 15, 911 (2014).

58. Kovárová, J. et al. Gluconeogenesis using glycerol as a substrate in bloodstream-form Trypanosoma brucei. PLoS Pathog. 14, (2018).

59. Haenni, S., Renggli, C. K., Fragoso, C. M., Oberle, M. & Roditi, I. The procyclin-associated genes of Trypanosoma brucei are not essential for cyclical transmission by tsetse. Mol. Biochem. Parasitol. 150, 144–156 (2006).

60. Lähnemann, D. et al. Eleven grand challenges in single-cell data science. Genome Biol. 21, (2020).

61. Qiu, P. Embracing the dropouts in single-cell RNA-seq analysis. Nat. Commun. 11, (2020).

62. Mou, T., Deng, W., Gu, F., Pawitan, Y. & Vu, T. N. Reproducibility of Methods to Detect Differentially Expressed Genes from Single-Cell RNA Sequencing. Front. Genet. 10, (2020).

63. Coley, A. F., Dodson, H. C., Morris, M. T. & Morris, J. C. Glycolysis in the African Trypanosome: Targeting Enzymes and Their Subcellular Compartments for Therapeutic Development. Mol. Biol. Int. 2011, 1–10 (2011).

64. Devlin, R. et al. Mapping replication dynamics in Trypanosoma brucei reveals a link with telomere transcription and antigenic variation. Elife 5, e12765 (2016).

65. Faria, J., Briggs, E. M., Black, J. A. & McCulloch, R. Emergence and adaptation of the cellular machinery directing antigenic variation in the African trypanosome. Curr. Opin. Microbiol. 70, 102209 (2022).

66. McCulloch, R. & Barry, J. D. A role for RAD51 and homologous recombination in Trypanosoma brucei antigenic variation. Genes Dev. 13, 2875–88 (1999).

67. Devlin, R., Marques, C. A. & McCulloch, R. Does DNA replication direct locus-specific recombination during host immune evasion by antigenic variation in the African trypanosome? Curr. Genet. 63, 441–449 (2017).

68. Saldivar, J. C. et al. An intrinsic S/G2 checkpoint enforced by ATR. Science (80-.). 361, 806–810 (2018).

69. Eykelenboom, J. K. et al. ATR activates the S-M checkpoint during unperturbed growth to ensure sufficient replication prior to mitotic onset. Cell Rep. 5, 1095–1107 (2013).

70. Marin, P. A. et al. ATR Kinase Is a Crucial Player Mediating the DNA Damage Response in Trypanosoma brucei. Front. Cell Dev. Biol. 8, 1651 (2020).

71. Black, J. A. et al. Trypanosoma brucei ATR Links DNA Damage Signaling during Antigenic Variation with Regulation of RNA Polymerase I-Transcribed Surface Antigens. Cell Rep. 30, 836-851.e5 (2020).

72. Glover, L., Marques, C. A., Suska, O. & Horn, D. Persistent DNA damage Foci and DNA replication with a broken chromosome in the African trypanosome. MBio 10, (2019).

73. Glover, L., McCulloch, R. & Horn, D. Sequence homology and microhomology dominate chromosomal double-strand break repair in African trypanosomes. Nucleic Acids Res. 36, 2608–18 (2008).

74. Li, Z. Regulation of the cell division cycle in Trypanosoma brucei. Eukaryot. Cell 11, 1180–90 (2012).

75. Wheeler, R. J., Gull, K. & Sunter, J. D. Coordination of the Cell Cycle in Trypanosomes. Annu. Rev. Microbiol. 73, 133–154 (2019).

76. Hammarton, T. C., Clark, J., Douglas, F., Boshart, M. & Mottram, J. C. Stage-specific differences in cell cycle control in Trypanosoma brucei revealed by RNA interference of a mitotic cyclin. J. Biol. Chem. (2003) doi:10.1074/jbc.M300813200.

77. Li, Z. & Wang, C. C. A PHO80-like cyclin and a B-type cyclin control the cell cycle of the procyclic form of Trypanosoma brucei. J. Biol. Chem. 278, 20652–20658 (2003).

78. Hayashi, H. & Akiyoshi, B. Degradation of cyclin B is critical for nuclear division in Trypanosoma brucei. Biol. Open 7, (2018).

79. Mugo, E. & Clayton, C. Expression of the RNA-binding protein RBP10 promotes the bloodstream-form differentiation state in Trypanosoma brucei. PLoS Pathog. (2017) doi:10.1371/journal.ppat.1006560.

80. Dejung, M. et al. Quantitative Proteomics Uncovers Novel Factors Involved in Developmental Differentiation of Trypanosoma brucei. PLoS Pathog. 12, e1005439 (2016).

81. Wurst, M. et al. Expression of the RNA recognition motif protein RBP10 promotes a bloodstream-form transcript pattern in Trypanosoma brucei. Mol. Microbiol. 83, 1048–1063 (2012).

82. Monnerat, S. et al. Identification and Functional Characterisation of CRK12:CYC9, a Novel Cyclin-Dependent Kinase (CDK)-Cyclin Complex in Trypanosoma brucei. PLoS One (2013) doi:10.1371/journal.pone.0067327.

83. Liu, Y., Hu, H. & Li, Z. The cooperative roles of PHO80-like cyclins in regulating the G1/S transition and posterior cytoskeletal morphogenesis in Trypanosoma brucei. Mol. Microbiol. 90, 130–146 (2013).

84. Li, Z. Regulation of the cell division cycle in Trypanosoma brucei. Eukaryot. Cell 11, 1180–90 (2012).

85. Mahdessian, D. et al. Spatiotemporal dissection of the cell cycle with single-cell proteogenomics. Nat. 2021 5907847 590, 649–654 (2021).

86. Clayton, C. Gene expression in Kinetoplastids. Curr. Opin. Microbiol. 32, 46–51 (2016).

87. Clayton, C. Regulation of gene expression in trypanosomatids: living with polycistronic transcription. Open Biol. 9, (2019).

88. Silvester, E., Ivens, A. & Matthews, K. R. A gene expression comparison of Trypanosoma brucei and Trypanosoma congolense in the bloodstream of the mammalian host reveals species-specific adaptations to density-dependent development. PLoS Negl. Trop. Dis. 12, e0006863 (2018).

89. Naguleswaran, A., Doiron, N. & Roditi, I. RNA-Seq analysis validates the use of culture-derived Trypanosoma brucei and provides new markers for mammalian and insect life-cycle stages. BMC Genomics 19, 227 (2018).

90. Gunasekera, K., Wüthrich, D., Braga-Lagache, S., Heller, M. & Ochsenreiter, T. Proteome remodelling during development from blood to insect-form Trypanosoma brucei quantified by SILAC and mass spectrometry. BMC Genomics 13, 556 (2012).

91. Hirumi, H. & Hirumi, K. Continuous cultivation of Trypanosoma brucei blood stream forms in a medium containing a low concentration of serum protein without feeder cell layers. J. Parasitol. 75, 985–9 (1989).

92. Brun, R. & Schönenberger. Cultivation and in vitro cloning or procyclic culture forms of Trypanosoma brucei in a semi-defined medium. Short communication. Acta Trop. 36, 289–292 (1979).

93. Rojas, F. et al. Oligopeptide Signaling through TbGPR89 Drives Trypanosome Quorum Sensing Article Oligopeptide Signaling through TbGPR89 Drives Trypanosome Quorum Sensing. Cell 176, 306-317.e16 (2019).

94. Vasquez, J. J., Hon, C. C., Vanselow, J. T., Schlosser, A. & Siegel, T. N. Comparative ribosome profiling reveals extensive translational complexity in different Trypanosoma brucei life cycle stages. Nucleic Acids Res. (2014) doi:10.1093/nar/gkt1386.

95. Ivens, A. C. et al. The genome of the kinetoplastid parasite, Leishmania major. Science 309, 436–442 (2005).

96. Amos, B. et al. VEuPathDB: the eukaryotic pathogen, vector and host bioinformatics resource center. Nucleic Acids Res. 50, D898–D911 (2022).

97. Love, M. I., Huber, W. & Anders, S. Moderated estimation of fold change and dispersion for RNA-seq data with DESeq2. Genome Biol. (2014) doi:10.1186/s13059-014-0550-8.

98. Andreatta, M., Berenstein, A. J. & Carmona, S. J. scGate: marker-based purification of cell types from heterogeneous single-cell RNA-seq datasets. Bioinformatics 38, 2642–2644 (2022).

99. Hao, Y. et al. Integrated analysis of multimodal single-cell data. Cell 184, 3573-3587.e29 (2021).

100. Finak, G., McDavid, A., … M. Y.-G. & 2015, undefined. MAST: a flexible statistical framework for assessing transcriptional changes and characterizing heterogeneity in single-cell RNA sequencing data. genomebiology.biomedcentral.com.

101. Haghverdi, L., Lun, A. T. L., Morgan, M. D. & Marioni, J. C. Batch effects in single-cell RNA-sequencing data are corrected by matching mutual nearest neighbors. Nat. Biotechnol. 2018 365 36, 421–427 (2018).

102. Benjamini, Y. & Hochberg, Y. Controlling the False Discovery Rate: A Practical and Powerful Approach to Multiple Testing. J. R. Stat. Soc. Ser. B 57, 289–300 (1995).

103. Berriman, M. The Genome of the African Trypanosome Trypanosoma brucei. Science (80-.). 309, 416–422 (2005).

104. Li, L., Stoeckert, C. J. & Roos, D. S. OrthoMCL: Identification of Ortholog Groups for Eukaryotic Genomes. Genome Res. 13, 2178 (2003).

105. Sievers, F. et al. Fast, scalable generation of high-quality protein multiple sequence alignments using Clustal Omega. Mol. Syst. Biol. 7, 539 (2011).

106. Lefort, V., Desper, R. & Gascuel, O. FastME 2.0: A Comprehensive, Accurate, and Fast Distance-Based Phylogeny Inference Program. Mol. Biol. Evol. 32, 2798–2800 (2015).

107. Beneke, T. & Gluenz, E. LeishGEdit: A Method for Rapid Gene Knockout and Tagging Using CRISPR-Cas9. Methods Mol. Biol. 1971, 189–210 (2019).

