## Supplementary figures for "Profiling the bloodstream form and procyclic form *Trypanosoma brucei* cell cycle using single cell transcriptomics"

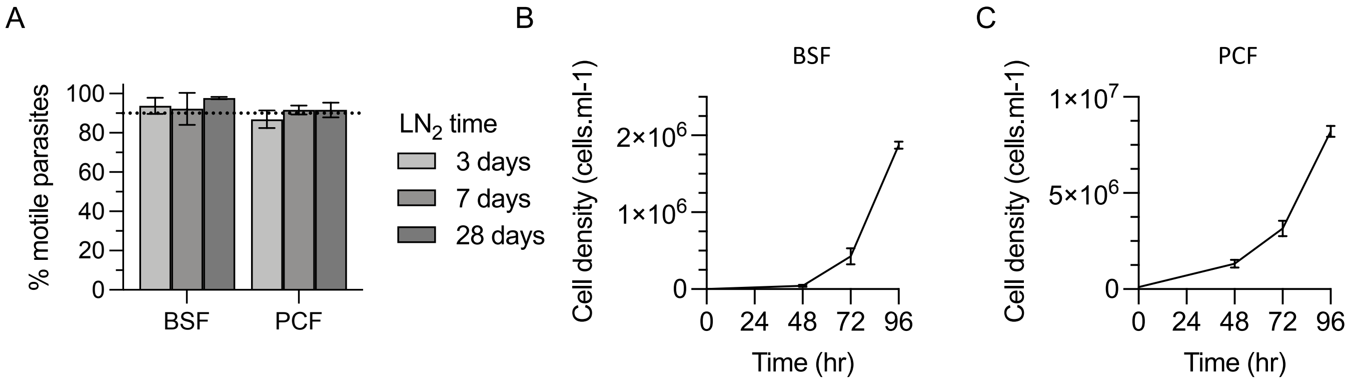


**Supplementary figure 1. Effect of cryopreservation on *T. brucei* viability**. The percentage of motile BSF and PCF parasites after preservation in $LN_2$ for 3, 7 or 28 days. Growth curves of BSFs (**b**) and PCFs (**c**) after being recovered from cryopreservation by the slow thawing protocol and returned to culture after 28 days of storage. Error bars show the SD from the mean of two independent replicates.

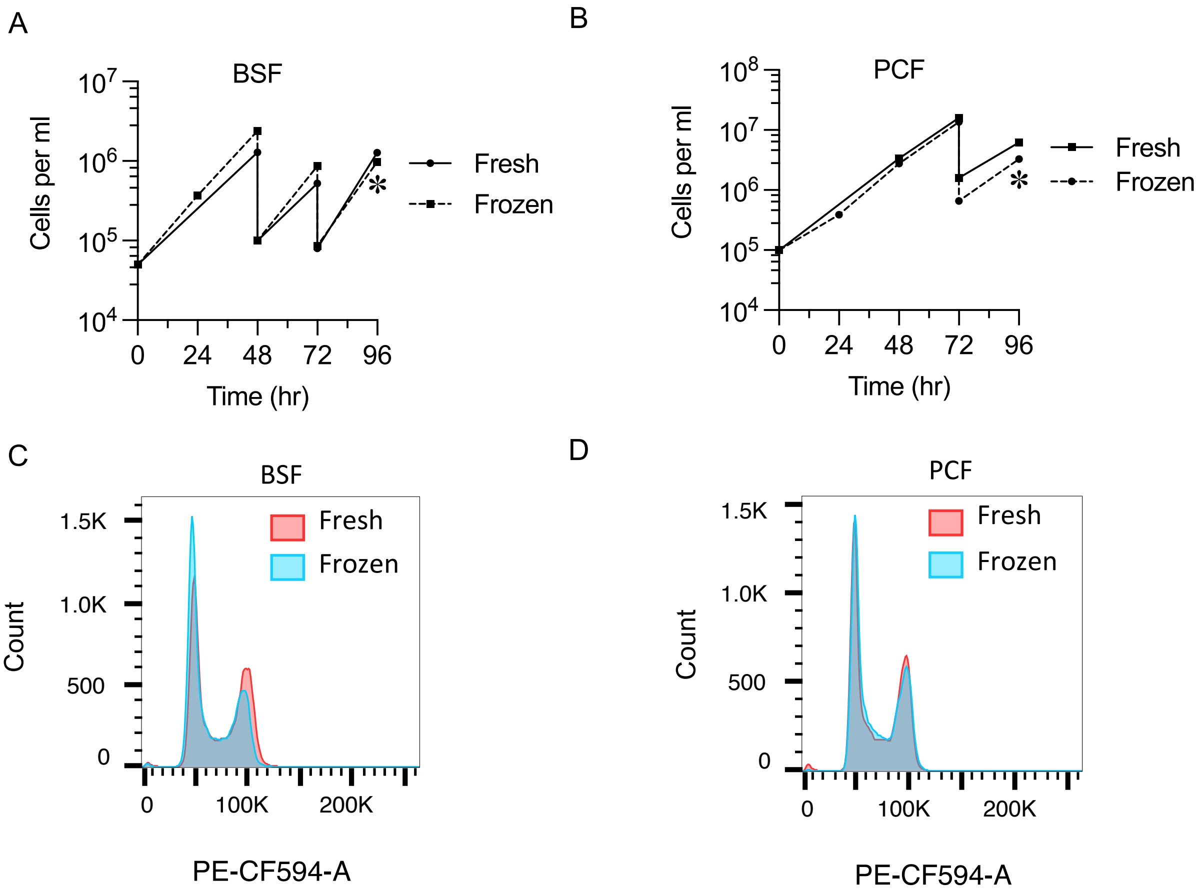


**Supplementary figure 2. Preparation of replicating BSF and PCF *T. brucei* prior to scRNA-seq or cryopreservation.** Growth of BSF (**a**) and PCF (**b**) *T. brucei* passaged over 4 days prior to immediate scRNA-seq preparation (Fresh, solid line) or cryopreservation (Frozen, dashed line) at the point indicated by an asterisk. Flow cytometry analysis of PI samples stained at the point indicated by asterisk for BSFs (**c**) and PCFs (**d**).

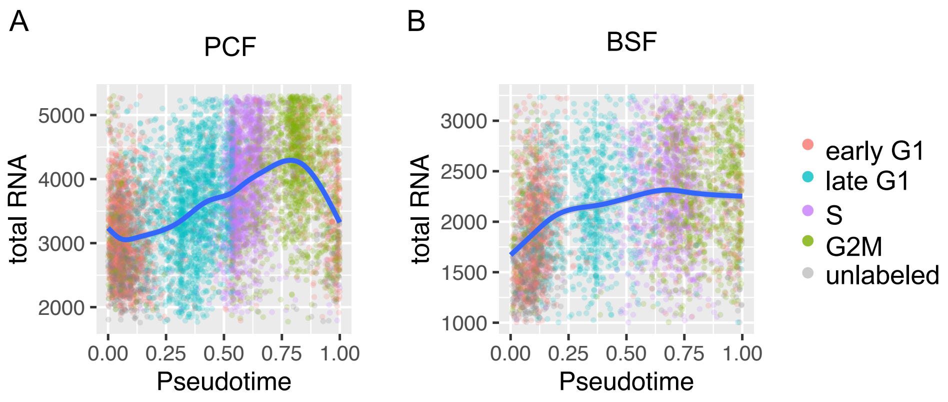


**Supplementary figure 3. Total RNA captured per cell across BSF and PCF cell cycle progression.** Total unique transcripts per cell is plotted on the y-axis (total RNA) over the inferred pseudotime (x-axis) for PCF (**a**) and BSF (**b**). Each cell is coloured by assigned cell cycle phase. Blue solid line indicates smoothed average RNA over pseudotime.

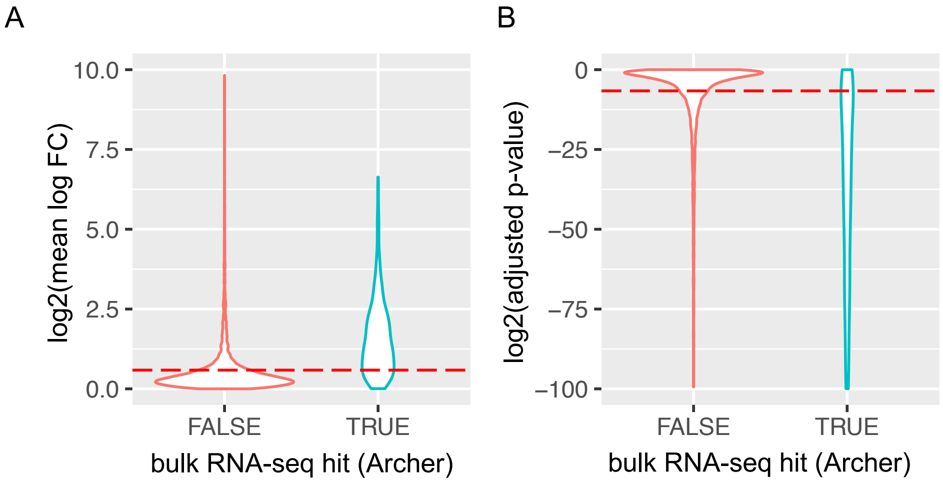


**Supplementary figure 4. Cell cycle regulated gene selection thresholds.** Violin plots indicate genes mean logFC over PCF pseudotime (**a**) and BH adjusted p-values (**b**) for genes (x-axis) grouped by whether they were previously identified at CCR using bulk RNA-seq (Archer *et al.* 2011). Red dashed lines indicates threshold used to define CCR genes in scRNA-seq analysis: FC > 1.5 and adjusted p-value < 0.01)

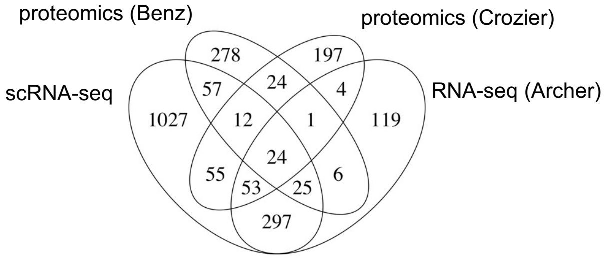


**Supplementary figure 5. Comparison of CCR genes selected in transcriptomic and proteomic studies.** Venn diagram indicates the number of common genes identified in the PCF scRNA-seq cell cycle analysis presented here, proteomic analysis from Crozier *et al.* 2018, proteomics analysis from Benz *et al* 2019 and bulk-RNA-seq from Archer *et al.* 2011. Genes were selected based on the same thresholds used in the original analysis of each study.

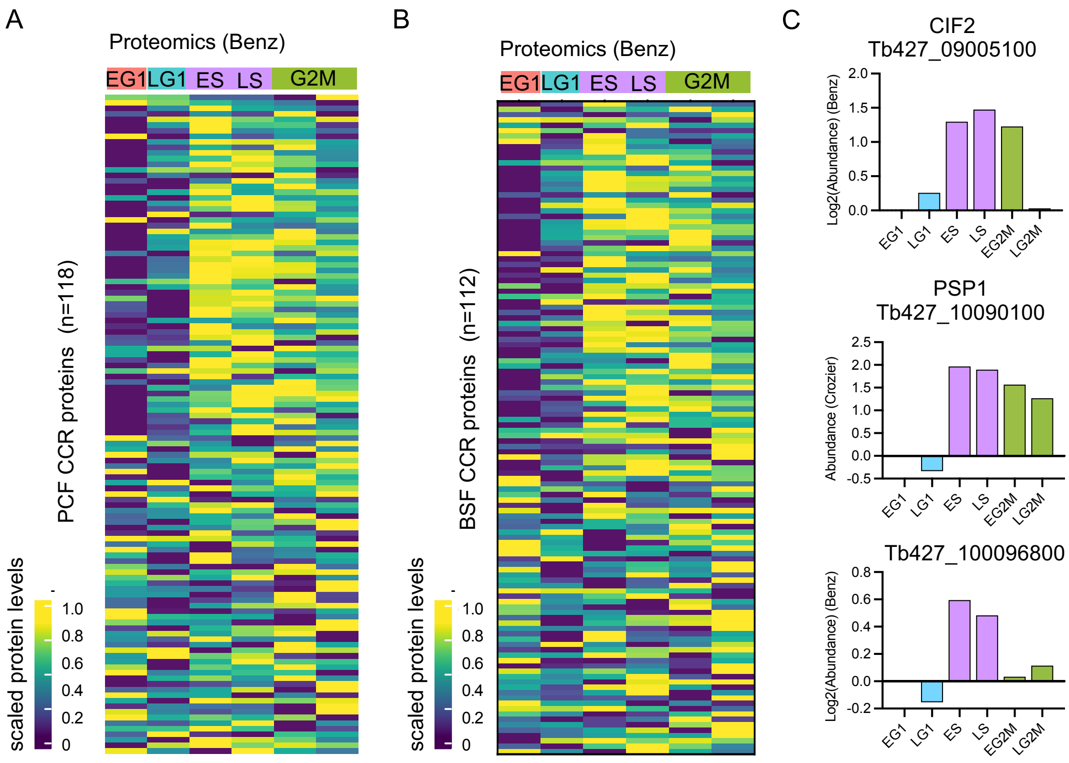


**Supplementary figure 6. Protein abundance levels across the PCF cell cycle as defined by Benz *et al*.** (**a**) Scaled protein abundance for 118 genes identified as CCR by Benz *et al* and scRNA-seq analysis of the PCF cell cycle, plotted in the same order as Fig. 2f. Time points are indicated in the top annotation, coloured by the most enriched cell cycle phase for each sample. (**b**) Scaled protein abundance for CCR 112 genes identified as CCR by Benz *et al* and scRNA-seq analysis of the BSF cell cycle, plotted in the same order as Fig. 3f. (**c**) Protein abundance from Benz *et al* study for the same genes as in Fig. 2j, previously identify as CCR by in all studies compared. Time point and colour of most enriched phase for each sample (x-axis). Protein abundance is scaled compared to early G1 (EG1), which is set to 0 for all genes.

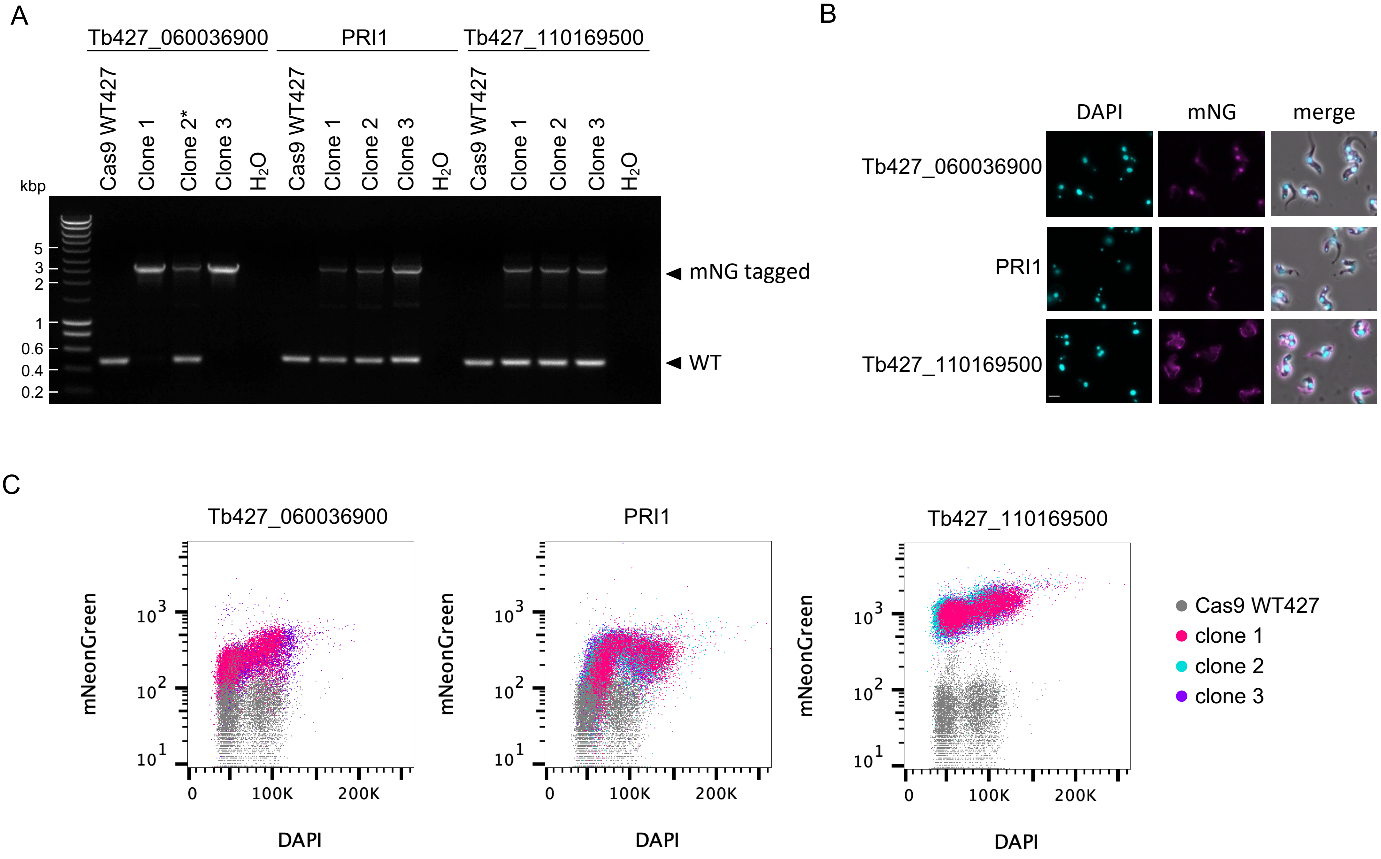


**Supplementary figure 7. Analysis of mNG tagged protein expression in BSFs. (a)** PCR detection of integrated mNG tag fragments for Tb427_060036900 (3,406 bp), PRI1/Tb427_080028700 (3,036 bp) and Tb427_110169500 (3,015 bp). Lower band indicates the WT without an inserted tag for Tb427_060036900 (484 bp), PRI1/Tb427_080028700 (498 bp) and Tb427_110169500 (477 bp). * indicates fluorescence could not be detected for this heterozygous Tb427_060036900::mNG clone. **(b)** Fluorescent microscopy imaging of mNeonGreen (mNG) tagged top CCR proteins. DAPI staining of DNA (cyan) and mNG fluorescence (magenta) are shown for the three genes as well as merged with DIC (merge). Scale bar = 10 µm **(c)** Scatter plots of parental Cas9 expressing WT427 BSFs (grey) and independently derived mNG tagged clones (pink, blue, purple) for three genes; Tb427_060036900, PRI1/Tb427_080028700 and Tb427_110169500). DAPI staining (x-axes) and mNG fluorescence (y-axes) were detected for 10,000 events per sample. Clone 2 is excluded from Tb427_060036900 as no fluorescence was detected.

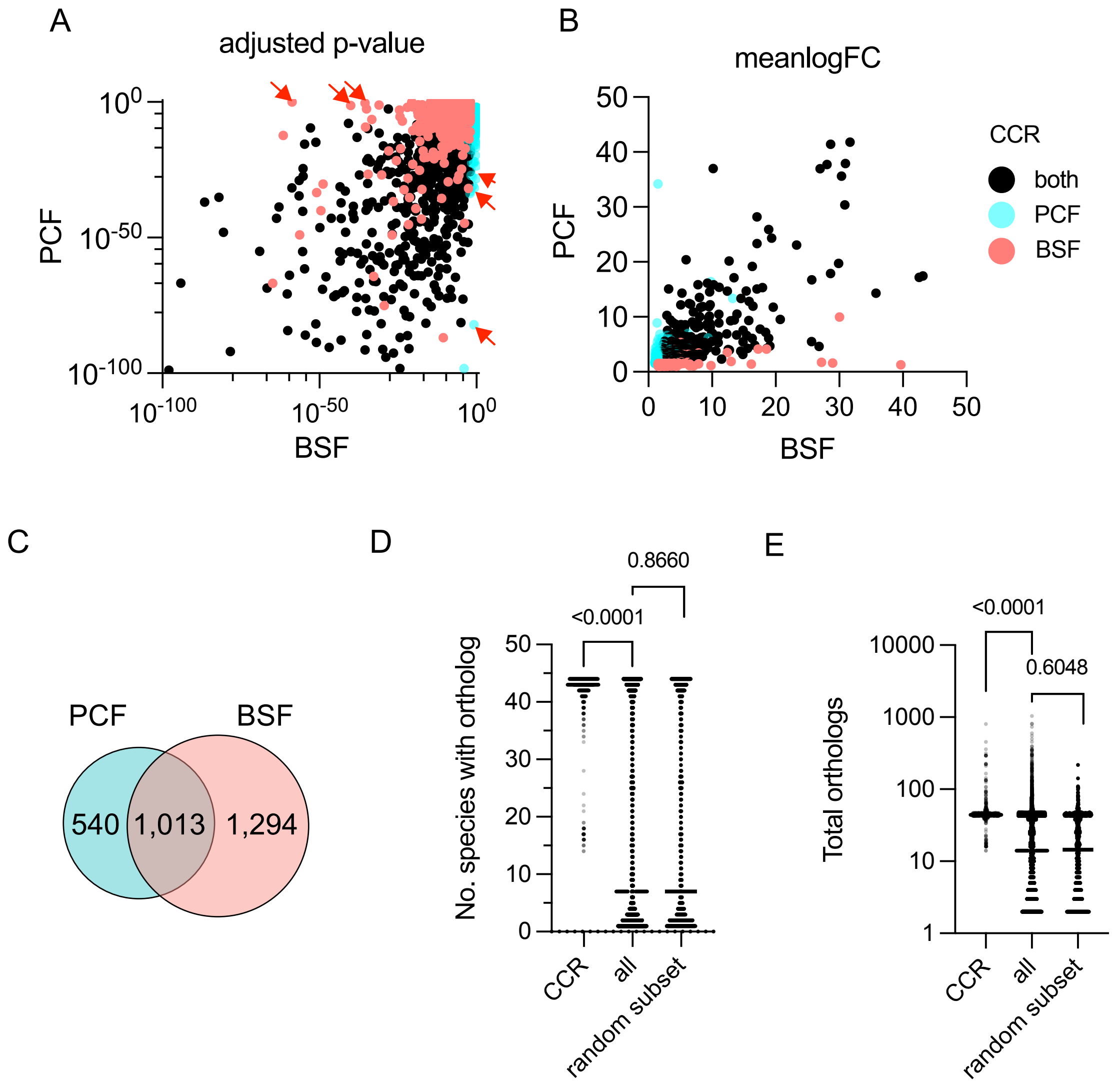


**Supplementary figure 8. Cell cycle regulated genes in PCF and BSF classified by adjusted p-value and FC.** (**a**) Scatter plot of genes classified as CCR in both life cycle forms (both, black), BSF only (red) or PCF only (blue). Genes are plotted by adjusted p-value for BSF (x-axis) and PCF (y-axis). (axis were limited to show data distribution removing 88 points). Red arrows indicate genes plotted in fig. 4e (**b**) CCR genes plotted by mean log FC in smoothed expression over pseudotime for BSFs (x-axis) and PCFs (y-axis). (18 points were removed due to limiting axes.) (**c**) Venn diagram demonstrating the overlap in PCF (blue) and BSF (red) CCR genes, defined by cut-offs in **a** and **b**. (**d**) Number of kinetoplastidea species reference genomes containing at least one ortholog of the common CCR genes, all genes from the WT427 reference genome, and a random subset of 1,000 genes. (**e**) Comparison of the total number of orthologs present across all 44 genomes for CCR genes, all genes and a random subset. p-values indicate the results of Mann-Whitney tests.

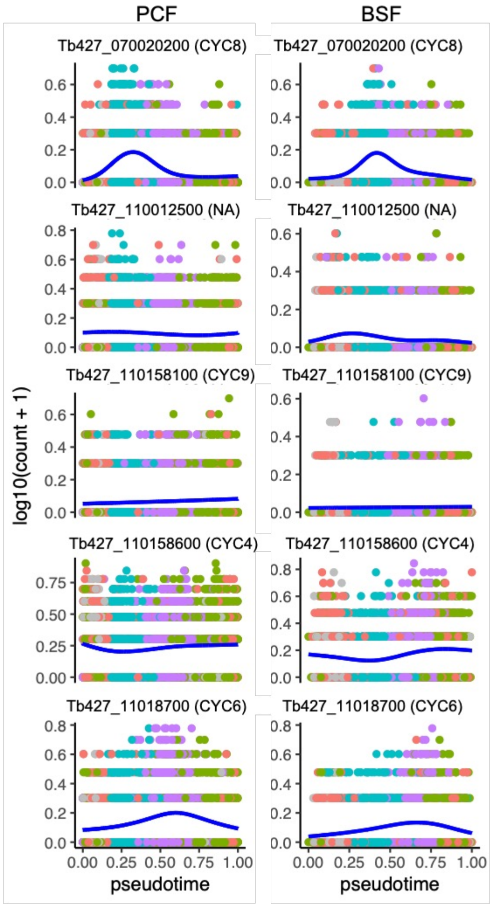


**Supplementary figure 9. Smoothed gene expression pattern of CCR cyclins.** Transcript levels of genes encoding known cyclins (CYC4, CYC6, CYC8 , CYC9) and one un-investigated cyclin domain containing gene (Tb427_110012500) found to be CCR in one or both forms. Counts per cell (y-axis) are plotted across PCF (left) and BSF (right) pseudotime (x-axis), coloured by phase. Blue line plots smoothed expression level across pseudotime.

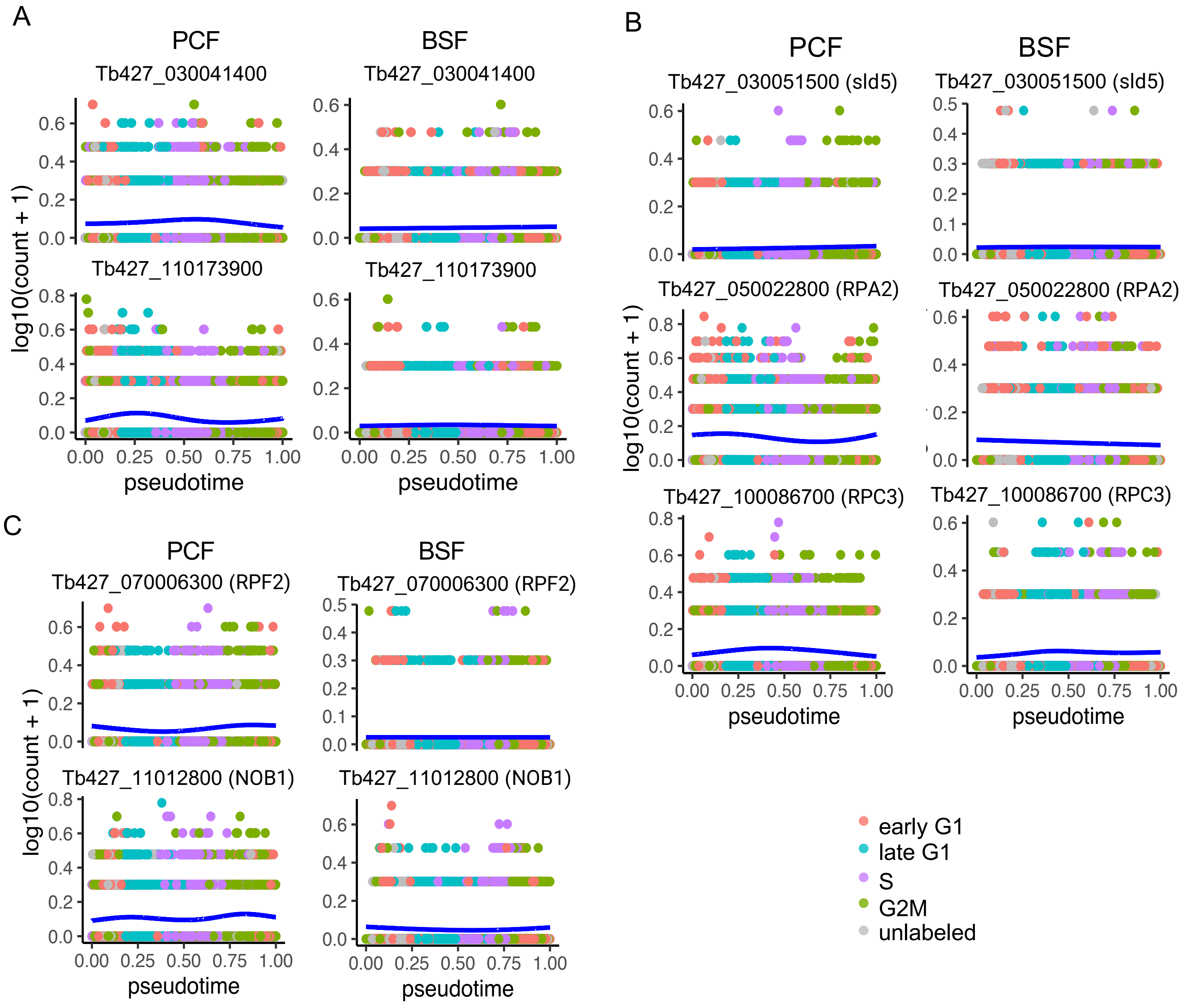


**Supplementary figure 10. Smoothed gene expression pattern of genes associated only with the PCF cell cycle.** Transcript levels of discussed example genes associated with GO terms “lipid metabolic process ” (**a**), " DNA replication ” (**b**) and “ribonucleoprotein complex biogenesis” (**c**). Counts per cell (y-axis) are plotted across PCF (left) and BSF (right) pseudotime (x-axis), coloured by phase. Blue line shows smoothed expression level across pseudotime.

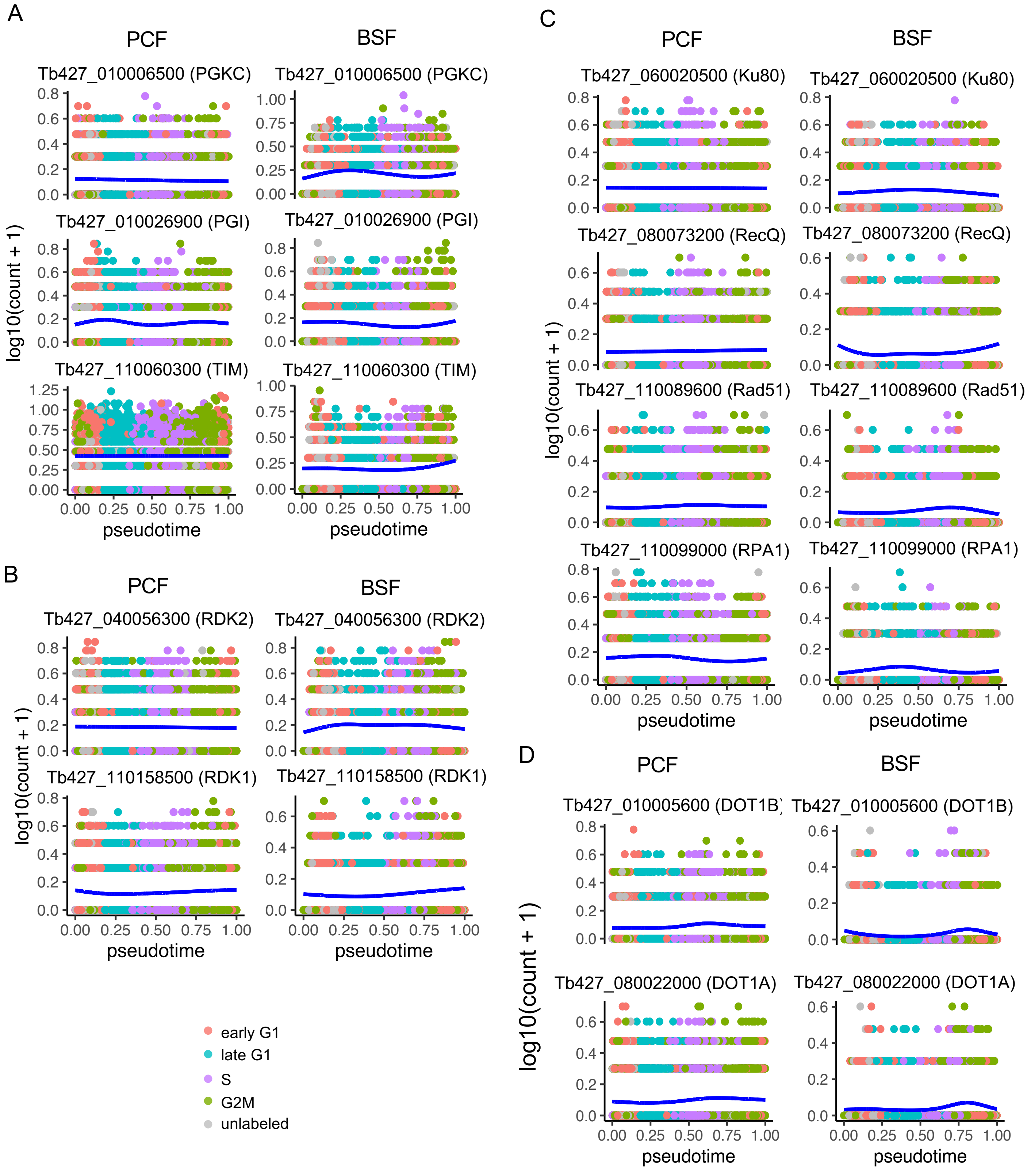


**Supplementary figure 11. Smoothed gene expression pattern of genes associated only with the BSF cell cycle** Transcript levels of discussed example genes for the glucose/gluconeogenesis pathway (**a**), and GO terms “phosphorylation” (**b**), “DNA recombination” (**c**) and "histone lysine methylation” (**d**). Counts per cell (y-axis) are plotted across PCF (left) and BSF (right) pseudotime (x-axis), coloured by phase. Blue line shows smoothed expression level across pseudotime.
